# Ribosomes modulate transcriptome abundance via generalized frameshift and out-of-frame mRNA decay

**DOI:** 10.1101/2024.03.12.584696

**Authors:** Yujie Zhang, Lilit Nersisyan, Eliska Fürst, Ioannis Alexopoulos, Susanne Huch, Claudio Bassot, Elena Garre, Per Sunnerhagen, Ilaria Piazza, Vicent Pelechano

**Affiliations:** SciLifeLab, Department of Microbiology, Tumor and Cell Biology, Karolinska Institutet, Solna, 171 65, Sweden; Armenian Bioinformatics Institute, Yerevan, Armenia; Institute of Molecular Biology, National Academy of Sciences of Armenia, Yerevan, Armenia; Max Delbrück Center for Molecular Medicine in the Helmholtz Association (MDC Berlin), Berlin, Germany; Department of Laboratory Medicine, Institute of Biomedicine, Sahlgrenska Academy, Sahlgrenska Center for Cancer Research, University of Gothenburg, 41390 Gothenburg, Sweden; Department of Chemistry and Molecular Biology, University of Gothenburg 40530 Gothenburg, Sweden

**Keywords:** NMD, mRNA decay, frameshift, codon optimality, out-of-frame

## Abstract

Cells need to adapt their transcriptome to quickly match cellular needs in changing environments. mRNA abundance can be controlled by altering both its synthesis and decay. Here we show how, in response to poor nutritional conditions, the bulk of the *S. cerevisiae* transcriptome undergoes -1 ribosome frameshifts and experiences an accelerated out-of-frame co-translational mRNA decay. Using RNA metabolic labelling, we demonstrate that in poor nutritional conditions, NMD-dependent degradation represents at least one third of the total mRNA decay. We further characterize this mechanism and identify low codon optimality as a key factor for ribosomes to induce out-of-frame mRNA decay. Finally, we show that this phenomenon is conserved from bacteria to humans. Our work provides evidence for a direct regulatory feedback mechanism coupling protein demand with the control of mRNA abundance to limit cellular growth and expands the functional role of mRNA quality control.

## Introduction

The modulation of gene expression in response to evolving environmental conditions is fundamental for cellular survival. This adaptive capability holds particular significance for unicellular organisms such as budding yeast or bacteria, as they rely on precise adjustment in gene expression to thrive amidst changing surroundings. mRNA abundance depends on the fine balance between mRNA synthesis and decay. mRNA decay controls the abundance of pre-existing mRNA molecules, modulates their availability for translation and facilitates rapid transcriptomic changes^1^. Defects in mRNA decay have been associated with multiple diseases ranging from neurodegeneration^2^ to viral infection^3^, underscoring its significance in controlling gene expression.

Multiple mechanisms control mRNA decay in response to environmental changes, for example, by destabilizing specific mRNAs in response to RNA binding proteins association^4^, by regulating the activity of proteins involved in mRNA decay^5,6^, or through co-translational mRNA decay^7,8,9^. A general process where the translation process modulates mRNA decay is the coupling between the demand of tRNAs by translating ribosomes and the available supply of charged tRNAs (codon optimality) which has been shown to regulate mRNA stability^10,11,12,13^. In addition to general processes controlling mRNA decay, multiple specialized mRNA surveillance pathways exist to ensure the elimination of faulty mRNAs and to facilitate ribosome recycling^12,14^. Classical examples include the <u>n</u>onsense <u>m</u>ediated <u>d</u>ecay pathway (NMD) associated with the elimination of transcripts containing <u>p</u>remature termination <u>c</u>odons (PTCs)^14,15,16^ or the <u>n</u>o-<u>g</u>o decay pathway (NGD) associated with the removal of mRNAs with stalled ribosomes^11^. Importantly, in addition to eliminating faulty transcripts, those pathways can also modulate the abundance of canonical mRNAs^1,14^. In general, NMD recognizes transcripts containing premature termination codons that can originate from genetic mutations, alternative splicing, or frameshifting events.

Frameshifts can regulate mRNA stability by causing premature translation termination and thus recruiting the NMD machinery. The frequency of spontaneous ribosome frameshifts is usually very low, as it requires the presence of a slippery sequence followed by a secondary structure element^17,18^. Frameshifts can also occur during ribosome translocation via tRNA slippage at the P-site while the A site is vacant^18^, especially associated with limitations in specific charged aa-tRNAs^19,20^. While ribosomal frameshifts have previously been linked to the degradation of specific transcripts, it is unclear if those events play a significant role in controlling global mRNA abundance, particularly when considering that most genes do not contain putative programmed ribosome frameshift (PRF) sites^21,22^.

We have previously shown that during co-translational mRNA decay, 5’-3’ exonucleases produce an *in vivo* toeprint of the position of the last (most 5’) trailing ribosome in yeast^8^ and bacteria^23^. Here, by investigating ribosome position associated with mRNA decay^8,24,25^, we discovered that the bulk of the *S. cerevisiae* transcriptome (∼77% of the degradation pool) undergoes -1 nt ribosome frameshifting in response to poor nutrient conditions. We characterize this process and identify both gene- and codon-specific features favouring frameshifting events. Next, we use genome-wide RNA metabolic labelling to demonstrate that in nutrient-poor conditions a sizable fraction of the transcriptome is degraded in an NMD-dependent manner.

We further characterized this mechanism and showed that low codon optimality, rather than the presence of PRF sites, is central to this process, and that amino acid supplementation can partially reverse this phenomenon. Next, we show that out-of-frame mRNA decay also contributes to changes in the proteome abundance. Surprisingly, this phenomenon is evolutionarily conserved and occurs not only in yeast and human cells but also in bacteria that lack canonical NMD machinery. Finally, we show that this new mechanism restricts cellular growth and conserves limiting resources under low nutrient conditions. We suggest that ribosome frameshifting followed by co-translational mRNA decay provides direct regulatory feedback coupling the demand of new proteins and the control of mRNA abundance encoding them.

## Results

### Study of co-translational mRNA decay reveals generalized -1 ribosome frameshifts

Sequencing the 5’P mRNA degradation intermediates naturally present in cells with 5PSeq provides the *in vivo* position of the last translating ribosomes^8^. 5PSeq is particularly well suited for investigating ribosome stalls associated with mRNA degradation, because it produces a toeprint of the subset of ribosomes engaged in co-translational mRNA decay, unlike ribosome profiling, which instead studies the bulk of ribosomes present in the cell^24,25^. In *S. cerevisiae*, ribosomes protect a region of 17 nt comprising the distance between the exposed 5’ phosphate (5’P) of an mRNA undergoing degradation and the ribosome A site. The ribosome protected region at the 5’ of mRNA has a constant size of 17 nt in different cellular conditions such as during oxidative stress, heat shock, cycloheximide treatment, or different growth media^8,25,26,27^. However, we serendipitously discovered that in very poor nutritional conditions the 17 nt ribosome protection pattern is displaced backward by 1 nt (−1 nt) (Fig 1A, Fig S1A). When cells are grown in <u>c</u>omplete <u>s</u>ynthetic <u>m</u>edia (CSM), this apparent -1 frameshift can be clearly observed in the body of the genes but not close to the start codon (Fig 1B-C). In fact, the -1 nt displaced frame (F0) only became predominant around 400 nt from the start (Fig 1C, middle). Surprisingly, we did not observe such alterations of the ribosome protection in other conditions with limited nutrients such as cells depleted for glucose, during early stationary phase or even using other synthetic defined media with slightly higher concentrations of nutrients (<u>s</u>ynthetic <u>c</u>omplete, SC media see methods) (Fig S1B-C). We observed a clear decrease of disomes in CSM media (Fig S1D), suggesting that ribosome collision is not driving this process.

**Figure 1.**
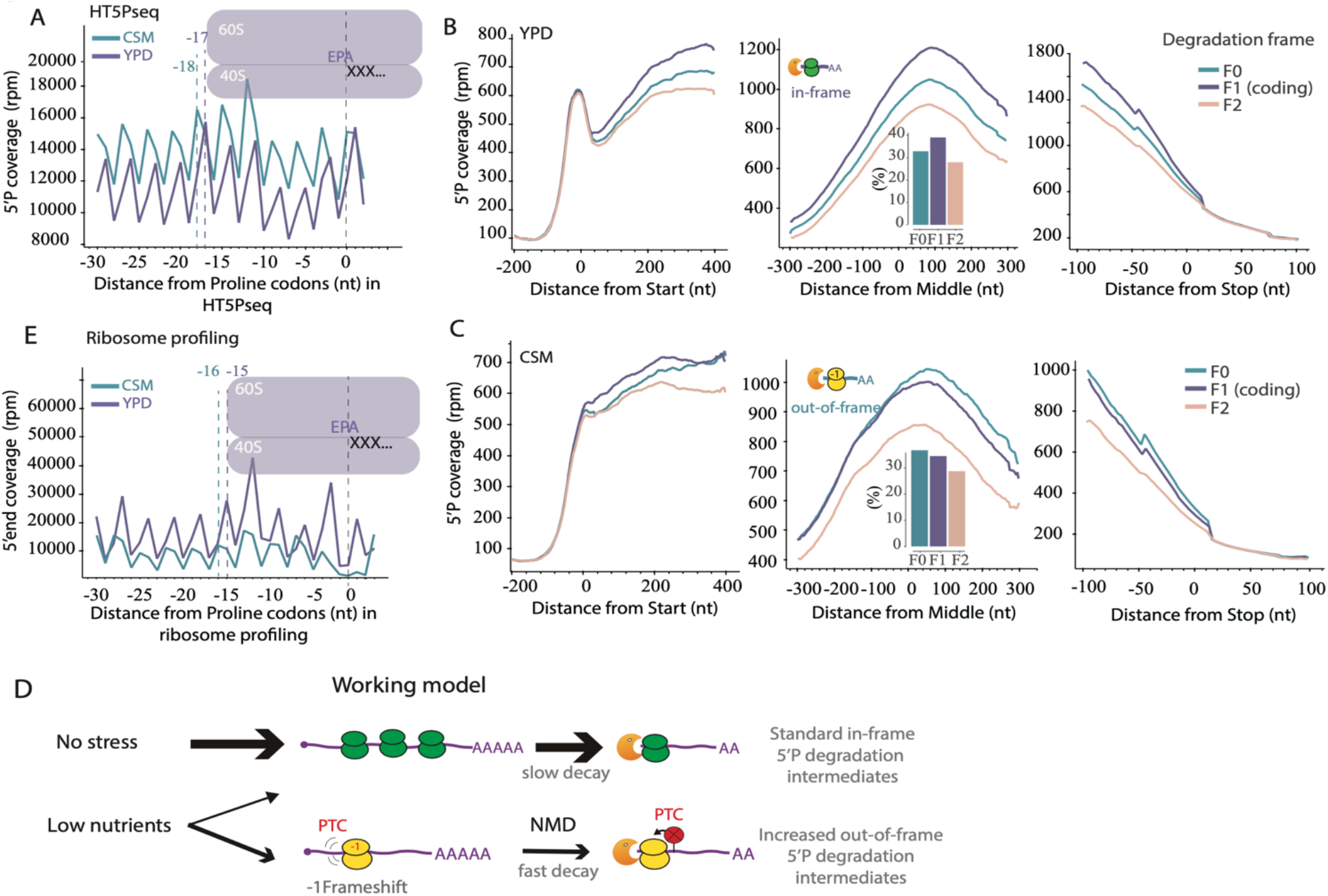
Study of co-translational mRNA decay reveals generalized -1 ribosome frameshifts. (A) Metagene analysis displaying the abundance of 5′P reads coverage for proline codons (CCG) in YPD (in purple) and in CSM (in green) using HT-5Pseq. Dotted lines at −17 and −18 correspond to the in-frame and *out-of-frame* 5′ end of protected ribosome located at the A-site, respectively. The same phenomenon can be seen for all codons, see Fig S1A. (B) Relative 5′P coverage for each frame in YPD from around the start codon (left), the middle of genes (middle) and the stop codon (right). A histogram displaying the relative coverage for each protection frame is shown. In 5PSeq, standard in-frame decay displays an increased coverage for F1, while a -1 nt frameshift will lead to a relative increase of coverage for F0. (C) As B, but for CSM. (D) Working model: in rich media conditions, *in-frame* mRNA degradation intermediates dominate the degradome. Under poor nutrient conditions, ribosomes will experience a higher frequency of -1 frameshifts. This will increase the proportion of *out-of-frame* mRNA degradation intermediates. mRNAs undergoing -1 frameshift would be likely degraded by NMD. (E) As A, but for ribosome profiling. Dotted lines at −15 and −16 correspond to the in-frame and *out-of-frame* 5′ end of protected ribosome located at the A-site as measured after *in vitro* RNase digestion, respectively.

We hypothesized two alternative scenarios that could explain why the apparent -1 frameshift does not occur proximal to the start codon: i) a gradual ribosome conformational change during late translation elongation leading to an altered *in vivo* 5’-3’ co-translational mRNA degradation pattern in the body of the genes; or ii) a massive increase of -1 ribosome frameshift frequency occurring during translation elongation and affecting the bulk of the transcriptome. As our data relies exclusively on the identification of *in vivo* 5’P boundaries we judged that an alternative ribosome conformation was unlikely. This was further supported by the fact that alternative protection sizes in ribosome profiling are caused by alternative 3’ boundaries depending on the *in vitro* RNase I accessibility to the ribosome A-site^28,29^ and not by changes in the 5’ side of the footprint.

To assess the likelihood of our second hypothesis, a massive increase of -1 ribosome frameshift frequency, we proposed a working model where, under poor nutrient conditions, ribosomes will undergo standard translation initiation, but will experience a -1 ribosome frameshift during translation elongation at an increased frequency (Fig 1D). At metagene level this would cause a -1 frameshift at some distance from the start codon, in agreement with our observations (Fig 1B-C). Frequent ribosome frameshifts will lead to the recognition of out-of-frame stop codons in the body of the genes and increased mRNA degradation via NMD. Reassuringly, we observed that putative *out-of-frame* stop codons increase 5PSeq ribosome footprints similar to those that we have previously described for canonical stop codons^25^ in CSM, but not in YPD (Fig S1E).

Our model suggests the existence of two populations of mRNA degradation intermediates: a canonical population with *in-frame* co-translational degradation and a second population subjected to frameshift-dependent NMD-enhanced degradation after -1 frameshift. At metagene level, and dependent on the relative importance of each pathway, we should observe canonical co-translational degradation profiles in the 5’ region of the genes (before the frameshift event) and potentially altered protection (−1 frameshift) in the body of the genes (after the frameshift). This scenario is consistent with our observations (Fig 1 and S1) and suggests that the bulk of the degradome of cells growing in poor media arises from a frameshift mediated degradation pathway.

Our working model also predicts that methods such as 5PSeq, focussing on transcripts undergoing decay^25^, should easily identify frameshift events associated with mRNAs subjected to increased decay. By contrast, in methods such as ribosome profiling, footprints derived from non-frameshifted ribosomes over stable mRNAs would likely mask frameshift events (Fig S1F-G). This would also explain why the dramatic phenomenon that we describe here has not been reported before. To test this scenario, we performed ribosome profiling in CSM conditions. We observed a clear increase of out-of-frame reads in CSM showing that ribosome frameshifting could also be detected using ribosome profiling, in a form of a -1 nt shoulder (Fig 1E and Fig S1F-H). However, the increase in out-of-frame protection measured by ribosome profiling was modest in comparison to the one measured by 5PSeq where out-of-frame ribosome protection clearly dominates (Fig 1A). Importantly, the observed shoulder agrees with previous observations showing that NMD-regulated transcripts tend to have a higher ratio of out-of-frame reads^30^. Taking all this together, our results confirm the existence of environmentally regulated genome-wide ribosome frameshift events enriched in mRNAs undergoing co-translational decay.

### Most genes experience environmentally induced -1 ribosome frameshifts

After showing the widespread existence of -1 frameshifts affecting the bulk of the transcriptome, we investigated the specificity of this process at a gene-specific level. We defined a simple metric to measure gene-specific frameshifts using the 3-nt periodicity associated with ribosome movement. For each gene we compared the in-frame 5PSeq sequencing coverage with respect to the coverage for a -1 frameshift (*i.e*., log_2_(F1/F0)). Using this metric only 212 genes (5.8 % of the 3645 analysed) present evidence for a -1 frameshift (log_2_(F1/F0) < 0) for exponentially growing cells in rich media (YPD) (Fig 2A, Fig S2A, Table S2A). In contrast, for exponentially growing cells in CSM -1 frameshift associated decay dominates the mRNA degradation in 2804 genes (77% of analysed) (Fig 2A, Fig S2A, Table S2A). To corroborate this, we performed the same analysis using ribosome profiling. In ribosome profiling the -1 frameshift is predominant for 34 genes in YPD (0.9%), and this number increases to 2326 genes in CSM (58%) (Fig S2B). This result confirms our previous observation showing that 5PSeq provides higher resolution to investigate ribosome frameshifts associated with mRNA decay (Fig 1A, 1E). Next, we investigated if this novel -1 frameshift phenomenon was specific to poor nutrition, or if it was induced also by other environmental challenges. To test these hypotheses, we investigated the effect of heat shock (30 min at 37 °C), oxidative stress (5 and 30 min after 0.2 mM H_2_O_2_ addition), and amino acid deprivation after growth in both rich and poor media (Fig 2B, S2C-J and Table S2B). Although the applied stressors differentially modulated the likelihood of frameshifts (as measured by the log_2_(F1/F0) ratio), it was clear that the used growth medium was the main driver of the phenotype (Fig 2B).

**Figure 2.**
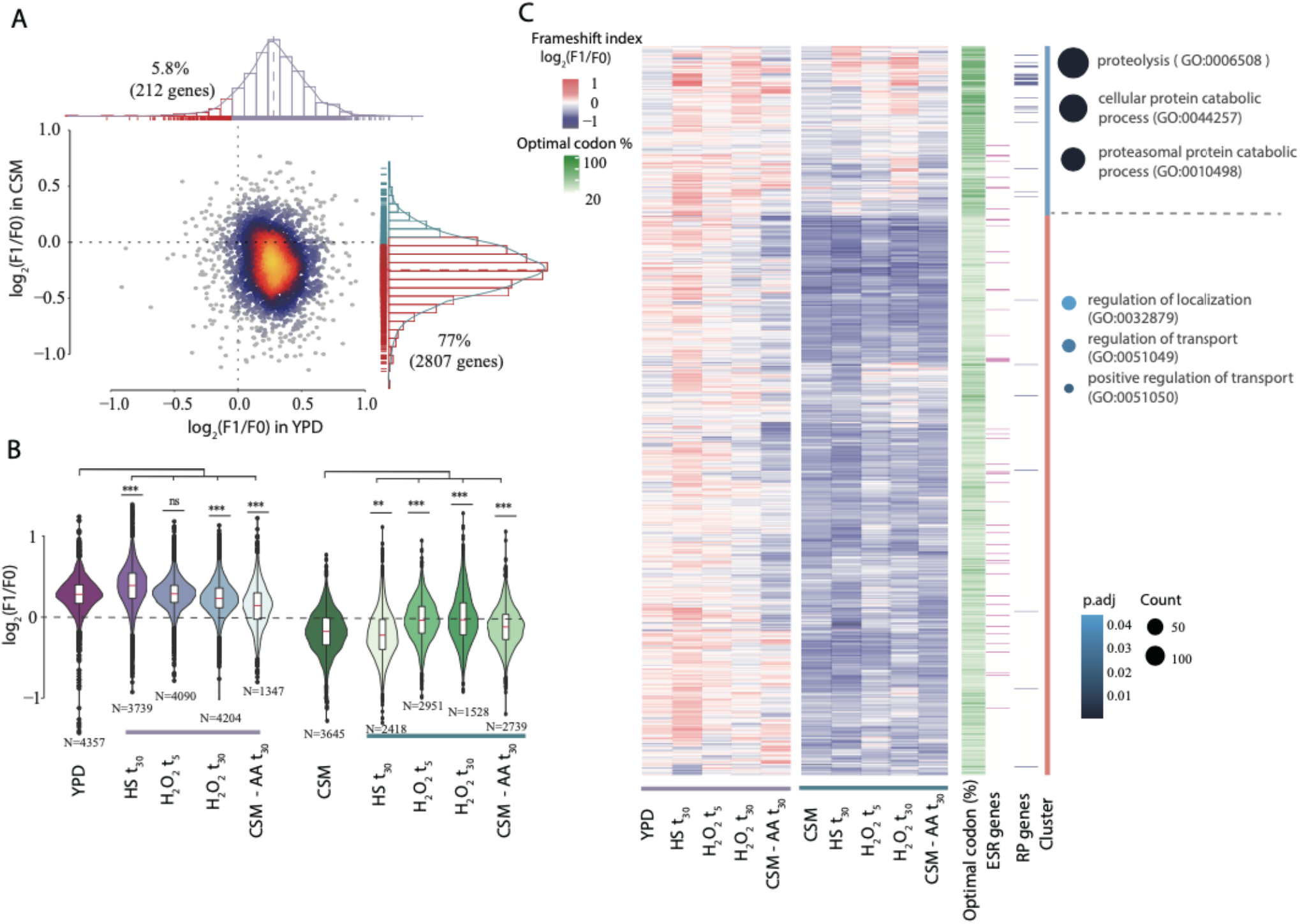
Gene-specific -1 ribosome frameshift are environmentally regulated. (A) Scatter plot comparing the frameshift index log_2_(F1/F0) for individual genes in rich (YPD, x-axis) and poor (CSM, y-axis) nutritional conditions. Histograms display the distribution of gene-specific frameshift index. Genes dominated by *out-of-frame* decay (log_2_(F1/F0) < 0) are highlighted in red, namely 212 genes in YPD and 2807 genes in CSM (5.8% and 77% respectively of total analysed genes). (B) Frameshift index distributions across tested conditions (as in B). The red line indicates the median gene frameshift index. Statistical analysis was performed using two-sided Wilcoxon rank-sum tests to its original condition (YPD and CSM) (*** P < 2.2 × 10^−16^; ** P < 6.3 × 10^−9^). Number of analysed genes are displayed. (C) Heatmap of frameshift index log_2_(F1/F0) for genes detected under all tested conditions starting from YPD (right) or CSM (left) clustered by k-means. Stresses include heat shock at 42°C for 30 mins, H_2_O_2_ (0.2 mM) exposure for 5 or 30 mins, and amino acid deprivation (transfer to CSM lacking amino acids) for 30 mins. Heatmap represents frameshift index log_2_(F1/F0), red to blue. Percentage of optimal codons for each gene are shown in green. Environmental stress response genes (ESR^31^) and ribosomal proteins genes are indicated in purple and blue, respectively. Frameshift index across all stress conditions were clustered using K-means (rightmost column). Gene ontology enrichments term for high and low frameshifted gene clusters are displayed (p-adj<0.05).

We used the 5PSeq data generated across the 10 tested growth conditions to investigate gene-specific -1 frameshifts (Fig 2C). Genes associated with regulation of RNA localization and intracellular protein transport presented a higher degree of frameshifting, while this was less pronounced in genes associated with proteolysis (Table S2C). Ribosome protein genes (RP) also had a relatively low tendency to experience an environment dependent -1 ribosome frameshift (Fig 2C and S2K). As we saw that stress conditions can modulate the level of frameshifting, we also investigated the behaviour of the Environmental Stress Response genes (ESR)^31^. However, we did not observe any clear association between the ESR genes and the frameshift events (Fig S2K).

Finally, we investigated if other gene-specific features could explain the observed differences in gene-specific frameshifting propensity. Factors such as gene length, 5’ UTR or 3’ UTR length did not affect the observed differences (Fig S2L-P). However, lower codon optimality^10^ and lower mRNA stability where clearly associated with the gene-specific frameshifting sensitivity (Fig 2C and S2M). We found the association between lower codon optimality and increased frameshifts particularly interesting, as it suggests a direct role of the ribosomes in this process. Additionally, our working model suggests that mRNAs experiencing a higher level of frameshifting events will be mainly degraded by an NMD-dependent pathway (Fig 1D), consistent with the fact that NMD-regulated transcripts tend to have lower codon optimality scores^30^.

### Global frameshifting in poor nutritional conditions promotes mRNA decay

A central prediction of our model is the co-existence of two alternative co-translational mRNA degradation pathways: a canonical *in-frame* decay and an accelerated frameshift-dependent *out-of-frame* one (Fig 1D). To estimate the fraction of the transcriptome degraded by each pathway, we took advantage of our generated 5PSeq data, which provides a snapshot of all mRNAs undergoing degradation. We assumed the fraction of out-of-frame transcripts in YPD to be nearly 0%, and then simulated a dataset with a 100% theoretical out-of-frame decay (shifting all YPD reads by -1 nt). By mixing reads of these 2 datasets, we estimated that at least 52% of the degradome originates from *out-of-frame* decay in CSM (Fig 3A-B, Table S3, see methods for details). This number is a very conservative estimate, as out-of-frame decay is likely not completely negligible in cells exponentially growing in YPD.

**Figure 3.**
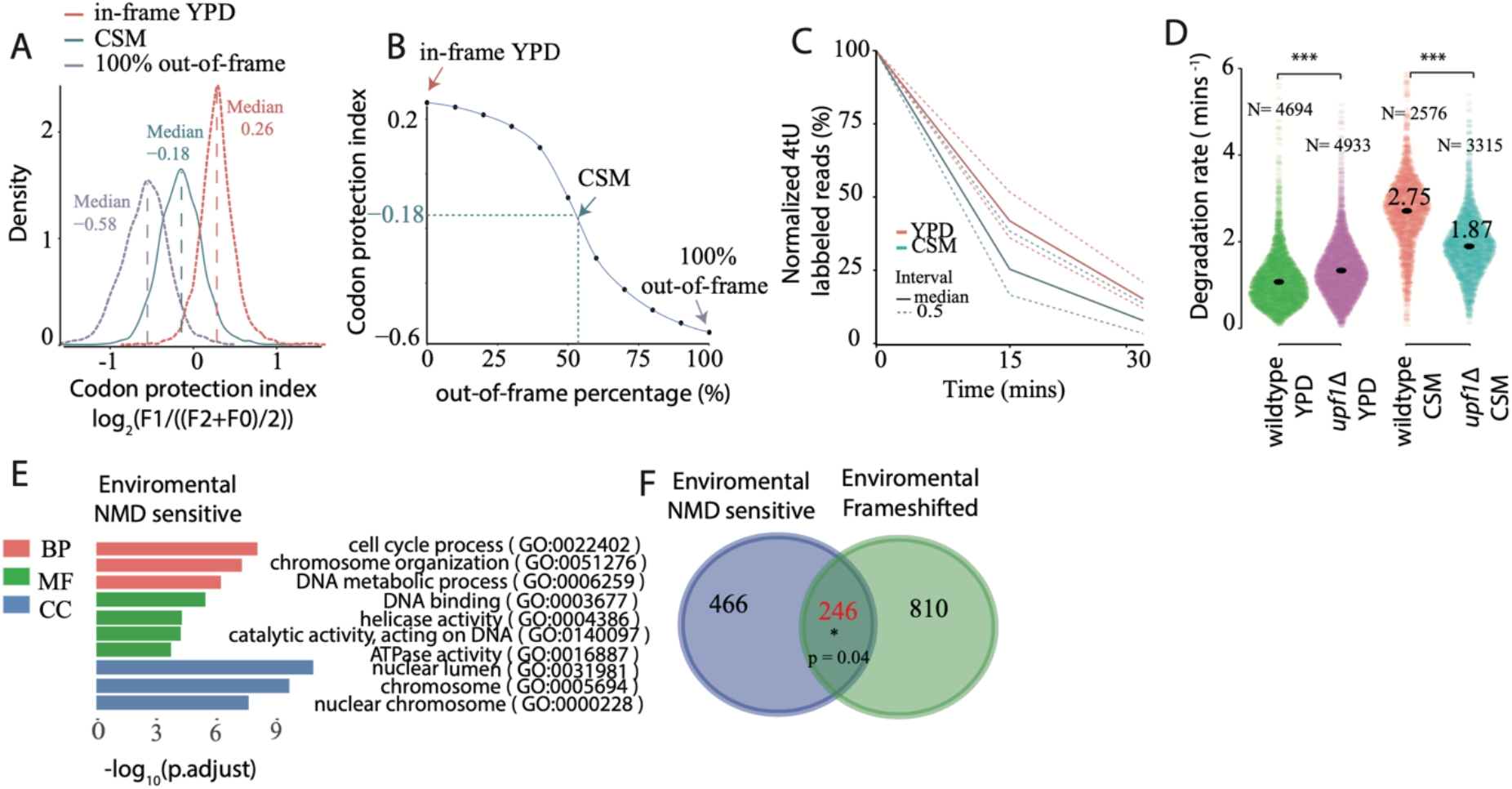
Ribosome frameshifts promotes mRNA degradation in poor nutritional conditions. (A) Distribution of gene-specific codon protection index for cells in YPD (median 0.26, red), in CSM (median -0.18, green) and simulated 100% *out-of-frame* decay (median -0.58, purple). Codon protection index was calculated as the ratio of the reads corresponding to the protected frame (F1) with respect to the average number of reads of the non-protected frames: log_2_(F1/((F2+F0)/2)). (B) Relationship between median codon protection index (y-axis) and percentage of *out-of-frame* reads (x-axis). Data was generated by mixing different ratios of 0% and 100% *out-of-frame* decay (see Methods). The green dotted line represents the median of codon protection index in CSM (−0.18) and corresponding estimated out-of-frame reads percentage (52 %). (C) Line plot displaying the average percentage of 4tU labelled reads after 15 and 30 mins 4tU pulse-chase. Data for YPD (pink) and CSM (green) are shown. Dotted lines provide the mean and median range. (D) Violin plot comparing the median degradation rate (mins^-1^) for wildtype (BY4741) and NMD mutant (*upf1Δ*) both in YPD and CSM. Only coding mRNAs with at least 20 total reads are considered for RNA turnover analysis. (E) Gene ontology terms for genes classified as environmentally dependent NMD sensitive genes (degradation rate (wildtype*/upf1Δ)*_CSM_ > 1.2 and (wildtype / *upf1Δ*)_YPD_ < 0.8) in three aspects (BP, biological process; CC, cellular component; MF, molecular function). Only top enrichments are shown (p-adjust < 0.01). Gene set universe (N= 2336) was set to the genes detected by SLAM-Seq across all conditions (Table S3A-C). (F) Overlap between genes subjected to environmentally dependent NMD decay and genes showing environmentally dependent frameshifts (log_2_(F1/F0) _YPD_ > 0.2 and log_2_(F1/F0) _CSM_ < -0.2). Statistical analysis was performed by hypergeometric analysis (k = 246, M = 712, N = 1257, s = 433) (Table S3D).

To test our model using an independent approach, we measured mRNA decay using pulse and chase RNA metabolic labelling followed by RNA-Sequencing (SLAM-Seq)^32^. This approach does not rely on the capture of transient mRNA degradation intermediates and measures disappearance of mRNAs, independent of the decay occurring co-translationally or not. We incubated cells for 60 minutes in a media containing 4-Thiouracil (4tU) and measured mRNA prior (t_0_) and after changing cells to a media without 4tU after 15 and 30 min in both YPD and CSM (see Methods for detail). Despite the known general association between faster cell growth and increased mRNA turnover in budding yeast^33^, we observed an increased mRNA decay (lower mRNA stability) in the slow growing cells in CSM (Fig 3C). This suggests that in CSM, in addition to the standard mRNA decay pathways associated with cell growth, another mechanism contributes to accelerated decay. As our previous results suggest that out*-of-frame* co-translational decay is associated with NMD, we compared the wild-type strain with an NMD-deficient strain (*upf1Δ*) (Fig 3D and S3B). In rich media, deletion of *UPF1* did not increase mRNA stability, and in fact led to a subtle increase of the mRNA degradation rate (suggesting potential adaptation of mRNA turnover in the *upf1Δ* strain). While growth rate was similar for both strains (Fig S3C). By contrast, in poor CSM conditions deletion of *UPF1* led to an increased mRNA stability (decrease of mRNA degradation rate). This confirms that in nutritionally poor conditions (in CSM), NMD is actively degrading a big fraction of the transcriptome. Using the generated mRNA metabolic data, we fitted the labelled mRNA abundance (normalized to total library size) across time to a non-linear decay model equation to calculate degradation rate for each condition^32^. We estimated that in CSM the median NMD-dependent degradation rate (measured from *upf1Δ*, 1.87 min^-1^) corresponds to at least 32% of total decay rate (measured from wildtype, 2.75 min^-1^), while non-NMD dependent degradation represent the remaining 68% (Fig 3D, Table S3A). Thus, even using conservative assumptions, under poor nutrition conditions at least one third of the transcriptome is degraded via environmentally induced ribosome frameshifts.

Next, we investigated the changes in gene-specific mRNA stability for cells with and without *UPF1* in rich and poor media. We classified those genes experiencing an increased NMD-dependent decay in CSM as environmental NMD-sensitive genes (*i.e*., the 712 genes where degradation rate (wt/*upf1Δ*)_CSM_ > 1.2 and (wt/*upf1Δ*)_YPD_ < 0.8). Those genes were enriched for cell cycle and chromosome organization (Fig 3D, Table S3B). Finally, we compared environmental NMD-sensitive genes with those experiencing environmentally induced frameshift (log_2_(F1/F0) _YPD_ > 0.2 and log_2_(F1/F0) _CSM_ < -0.2), see in Methods) and found a significant overlap (p-value = 0.04, Fig 3F, Table S3C). Taking all these together, we concluded that *out-of-frame* NMD-dependent degradation is responsible for the degradation of an important fraction of the transcriptome in poor nutritional conditions.

### Codon optimality controls environmentally regulated - 1 frameshifts and out-of-frame mRNA decay

After confirming the genome-wide nature of the environmentally regulated -1 frameshifts and their consequences for mRNA stability, we focused on their mechanisms. First, we investigated whether known mRNA slippery sequences^21,22^ caused the frameshifts. We reasoned that ribosome protection frames should be different before and after the ribosomes encounters those regions if it was the case. However, we did not observe any evidence for enhanced frameshifting surrounding slippery sequences (Fig S4A).

Since we observed massive frameshifts during poor nutrient conditions, the ribosome frameshifting could be due to limited availability of charged tRNAs. This is a phenomenon previously described for specific codons or mRNAs^34,35^ named “hungry codon” frameshift^19,20^. To test if a similar mechanism could be operating at genome-wide scale, we measured the degree of frameshift associated with each codon. We computed a relative protection frame for each codon (*i.e*., comparing 5PSeq coverage at -17 nt (F1, *in-frame*) and -18 nt (F0, *out-of-frame*) from the A-site; Fig 4A). Next, we compared the change in protection frame between YPD and CSM and defined a frameshift index such that codons more likely to induce a frameshift will present a higher value (*i.e*., log_2_(F1/F0)_YPD_ – log_2_(F1/F0)_CSM_). This analysis revealed that codons with lower optimality were more likely to engage in frameshifting in poor nutrient conditions. To further test this, we performed the same analysis using ribosome profiling data, which showed a similar trend, albeit with less sensitivity (Fig 4B and S4B). To investigate the impact of codon abundance at the gene-specific level, we compared the frameshift index distributions between YPD and CSM (Fig S4C) and focused on genes with high frameshift values (log_2_(F1/F0)_YPD_ > 0.2 and log_2_(F1/F0)_CSM_ < -0.2). We then calculated the Pearson correlation coefficient between the measured frameshift change (*i.e*., log_2_(F1/F0)_YPD_ - log_2_(F1/F0)_CSM_) and codon occurrence among these genes (Fig S4D). Consistent with our expectations, we found a positive correlation between a high frameshift index and the abundance of non-optimal codons, while low frameshift values were associated with a greater prevalence of optimal codons. All these results agree with our previous observation that mRNAs with lower codon optimality tend to display more frameshifts in CSM conditions (Fig 2B and S2O). Finally, we reasoned that in those conditions where a rare codon was in the A-site, and a -1 ribosome frameshift could lead to the incorporation of a common tRNA, the likelihood of frameshifting could increase. However, the nature of the incorporated codons after the frameshift played a minor role in this phenomenon (Fig 4C and S4E).

**Figure 4.**
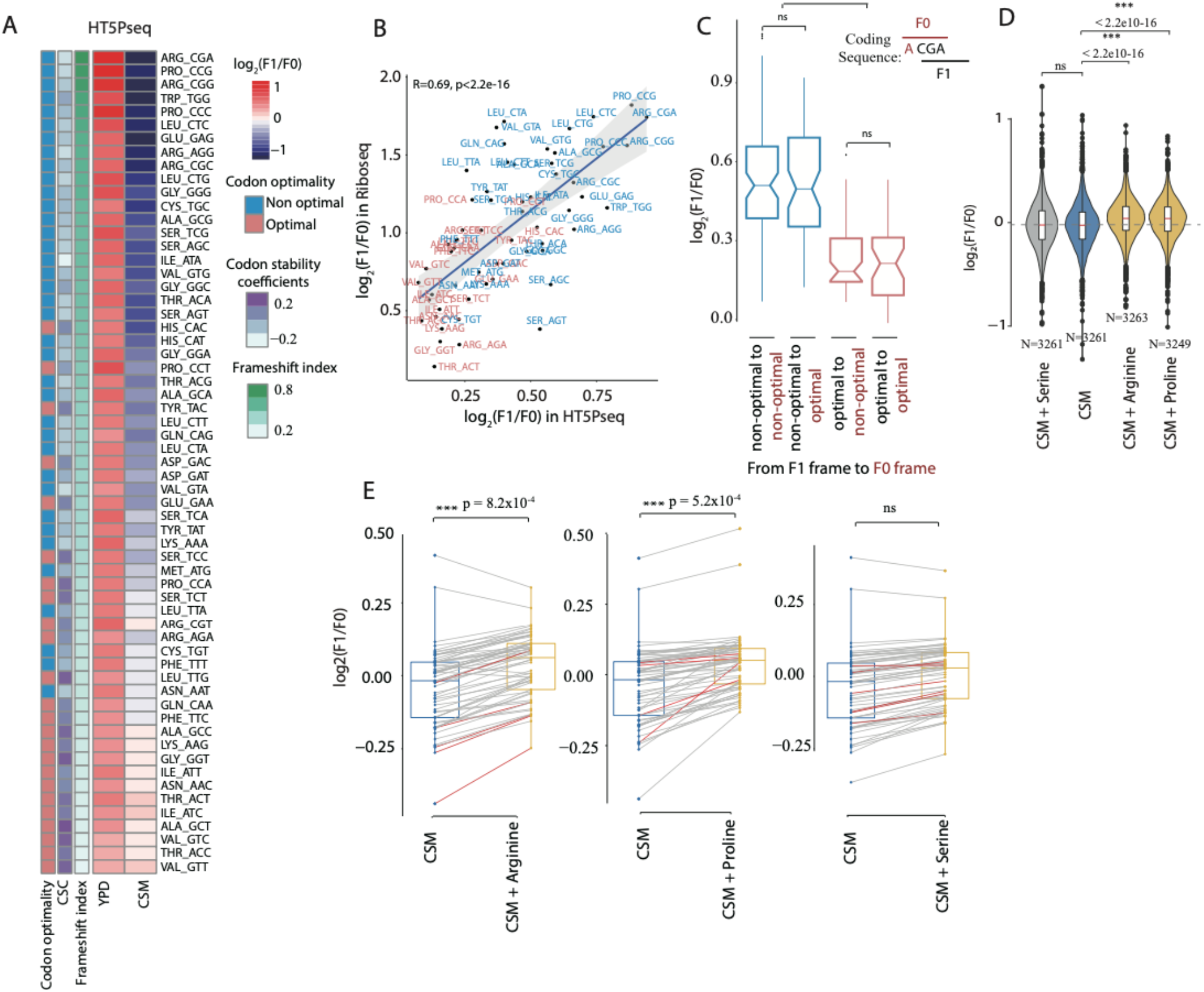
Codon optimality controls out-of-frame mRNA decay. (A) Heatmap comparing 5PSeq coverage at log_2_ ratio of -17 nt (F1) and -18 nt (F0) relative to the A-site of codons in YPD and CSM. The codons are ordered by the differences of frameshift index between YPD and CSM (*i.e*., log_2_(F1/F0)_YPD_ – log_2_(F1/F0)_CSM_). Codon stability coefficients (CSC) and Codon optimality from Presnyak *et al*.^10^ (B) Scatterplot comparing frameshift index using HT5PSeq and ribosome profiling. Spearman correlation is shown. Optimal codons and non-optimal codons are represented in red and blue, respectively. (C) Boxplot comparing the frameshift index among different combinations. The four combinations include: 1) non-optimal codons at both 17 nt (F1) and -18 nt (F0); 2) non-optimal codons at F1 and optimal codons at F0; 3) optimal codons at F1 and non-optimal codons at F0; 4) optimal codons at both F1 and F0. Non-optimal codons at F1 (in frame) are represented by blue and optimal codons at F1 are represented by red. Statistical analysis was performed using two-sided Wilcoxon rank-sum tests. *** P < 2.2 × 10^−16^. (D) Metagene plot showing the frameshift index change after amino acid supplementation. Serine, arginine or proline were added to reach the comparable concentration as SC medium. Statistical analysis was performed using two-sided Wilcoxon rank-sum tests. *** P < 2.2 × 10^−16^; ns, P = 0.58; Number of analysed genes are displayed. (E) Boxplot comparing frameshift index change per codons between CSM and the respective amino acid additions. The codons highlighted in red corresponding to respective amino acid (arginine, CGG/CGA/CGC/AGA; proline, CCT/CCC/CCA/CCG; serine, TCT/TCC/TCA/TCG/ AGT/AGC). Statistical analysis was performed using two-sided t tests.

To experimentally validate if the continued low amino acid availability for cells growing exponentially in CSM was a key driver of the observed phenomenon, we raised the final concentration of selected amino acids in CSM to the one present in SC media (see Methods). We first increased the concentration of amino acids whose codons are associated with higher frequency of frameshifts: arginine and proline (from 50 to 85.6 mg/L and 0 to 85.6 mg/L respectively). Reassuringly, both decreased the observed genome-wide frameshift events with respect to CSM (p-value <2.2·10^−16^) (Fig 4D). However, those changes were not restricted to the supplemented amino acids (Fig 4E and S4F) as could be expected by the fact that metabolic processes will enable their interconversion. Next, we increased serine concentration (from 0 to 85.6 mg/L), for which codons are in general associated with lower frameshift frequency (Fig 4A-B). As expected, serine addition had the least effect on the change of the log_2_(F1/F0) metric (Fig S4G). This shows that long term limitation in amino acid availability contributes to the appearance of genome-wide frameshifts and accelerates mRNA degradation.

### Long-term amino acid limitation rewires cellular proteome downstream of ribosome frameshifting

To further characterise the cellular state during conditions with high frameshifting, we analysed the proteome of cells grown in CSM and YPD. We identified 487 proteins differentially expressed (Q-value < 0.01), being 315 proteins significantly upregulated in CSM (log_2_FC > 1) and 172 proteins significantly downregulated (log_2_FC < -1) (Fig 5A, Table S5). Upregulated proteins involved mainly the proteasome complex, amino acid metabolic process and ergosterol biosynthetic process (Fig 5B and S5A), supporting that cells grown in CSM have a limited availability of amino acids in the media and thus need to upregulate the amino acid biosynthesis. While downregulated proteins were associated with components such as ribosomes, mitochondrial ribosome and nucleosome, as expected for a slower cellular growth in CSM.

**Figure 5.**
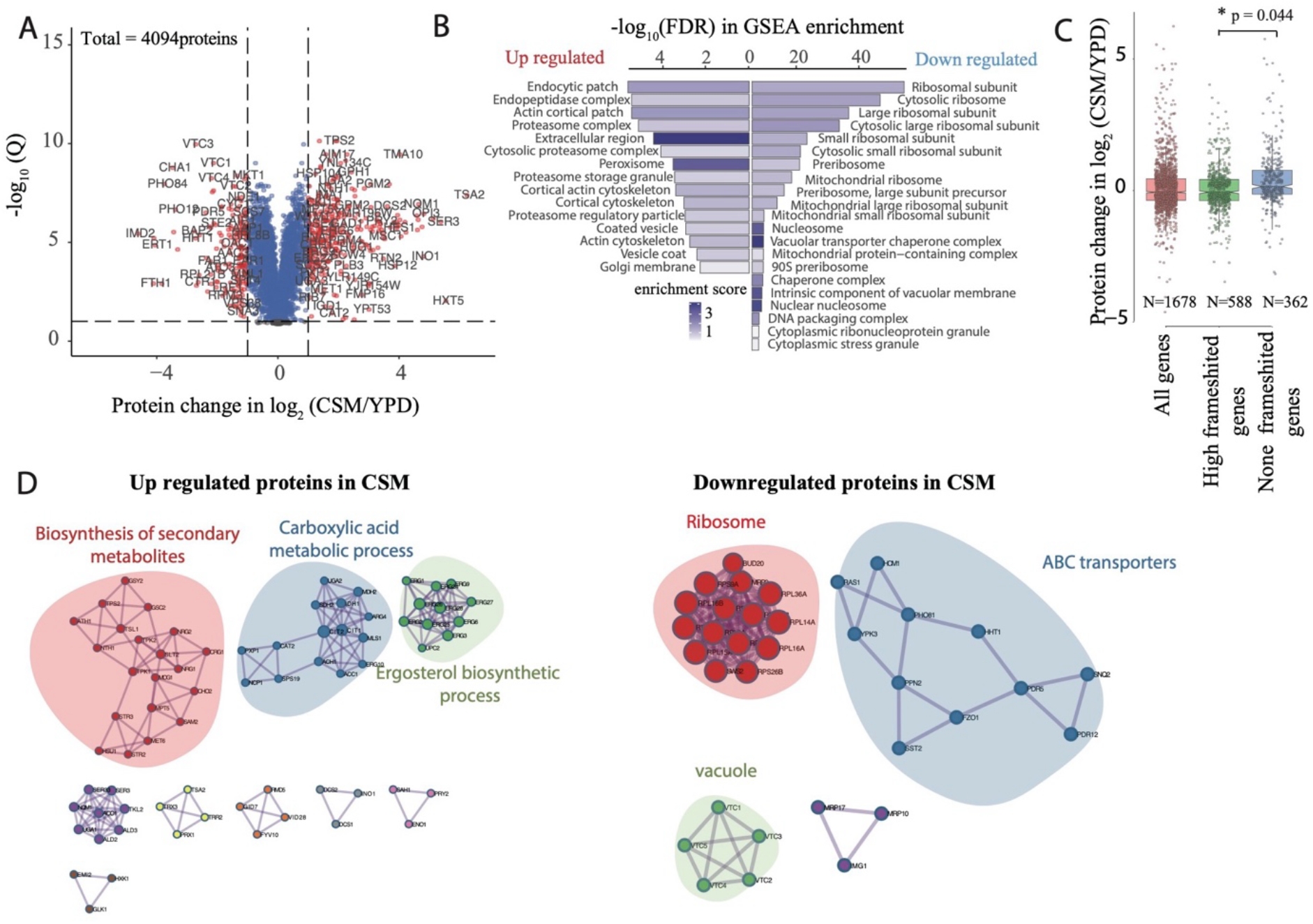
Long-term amino acid limitation rewires cellular proteome. (A) Volcano plot comparing protein abundance change between YPD and CSM in log_2_ fold change (x-axis) versus the statistical significance (−log_10_Q). We used a threshold of Q-value < 0.01 and log_2_FC >1 or log_2_FC < -1 to define 315 up-regulated and 172 down-regulated proteins. Only proteins with at least 2 peptides were considered for analysis (4094 proteins in total). (B) Gene set enrichment analysis (GSEA) for differential proteins abundance ranking according to log_2_FC with ClusterProfiler using Fisher’s exact test with p adjusted value < 0.05. (C) Boxplot comparing protein abundance levels for genes that are also detected in 5Pseq. Genes categorized as all genes (N = 1440), high frameshifted genes (log2(F1/F0) _YPD_ > 0.2 and log_2_(F1/F0) _CSM_ < -0.2, N =496 genes) and not frameshifted genes (log2(F1/F0) _YPD & CSM_ > 0, N = 319 genes). Statistical analysis is performed using Wilcoxon rank-sum tests with p value < 0.05. (D) Up-regulated and down-regulated proteins highlighting their protein-protein interactions. Analysis was performed using performed metascape v3.5. Only statistically significant enrichments are shown (p-adj < 0.05).

Next, we investigated if the observed frameshifts, in addition to modulating mRNA stability, also led to changes in protein abundance. We compared protein abundance change for those genes experiencing a relatively high level of frameshift (i.e., 588 genes where log_2_(F1/F0)_YPD_ > 0.2 and log_2_(F1/F0)_CSM_ < -0.2 and proteins are detected (Q-value < 0.01)) with those without frameshifts (362 genes where log_2_(F1/F0 _YPD & CSM_) > 0). We observed that protein abundance for genes with a higher frameshifting decreases more drastically in CSM than the relative protein abundance for those genes without frameshifts (Fig 5C). This confirms that, in addition to regulating the mRNA stability, this process also leads to differential protein abundance.

### Accelerated *out-of-frame* co-translational decay of mRNA is evolutionary conserved

Having characterized environmentally induced *out-of-frame* co-translational mRNA decay in *S. cerevisiae*, we explored the evolutionary conservation of this process. We reasoned those unicellular organisms, which are more exposed to the environment and need to adapt faster to changing conditions, should in principle be more susceptible to this phenomenon. As we have recently shown that 5’-3’ co-translational mRNA decay is also frequent in bacteria^23^, we investigated this process in *Lactobacillus plantarum* where the 5’-3’ exonuclease RNase J generates an *in vivo* toeprinting of the bacterial ribosome (14 nt from the A-site). We reanalysed our previous data^9^ where *L. plantarum* was grown exponentially in rich (MRS broth) and transferred to low-nutrient media (0.5xLB media) for 15 minutes. Despite the short exposure to low-nutrient conditions, we observed a significant increase for -1 frameshift (log_2_(F1/F0) <0) events in 6% of genes, up from 2% (48 genes) under rich conditions (Fig6A, Table S6).

To further test the conservation of this phenomenon at longer times, we investigated the 5’P degradome for *Bacillus subtilis*. Specifically, we compared the co-translational ribosome protection pattern for *B. subtilis* exponentially growing in rich (LB) and in poor media (Minimal growth medium, see in Methods). Reassuringly we observed a similar pattern where only 83 genes (3 % of measured) present evidence for -1 frameshifts in rich media, but this number increased to 336 genes (12%) in poor medium (Fig 6B). This pattern also held true when we checked *B. subtilis* during early stationary phase (after 24 hours growth in LB), where the number of genes dominated by -1 frameshifts increased to 17% of measured genes (Fig S6A). This shows that environmentally dependent *out-of-frame* co-translational mRNA decay that we described in budding yeast (Fig S4C) is also common in prokaryotes even in the absence of NMD. To investigate if the -1 frameshift followed a similar mechanism to the one studied in eukaryotes, we analysed the codon compositions of genes with varying susceptibility to frameshifting. Our findings, similar to those in yeast (Fig S4C), show a clear positive correlation between a high frameshift change (*i.e*., log_2_(F1/F0)_control_ – log_2_(F1/F0)_treatment_) and a high occurrence of non-optimal codons among genes with frameshifts (log2(F1/F0)_control_ > 0 and log2(F1/F0)_treatment_ < 0) and vice versa (Fig 6C-D and S6B-C). This analysis demonstrates, that also in prokaryotes, mRNAs more prone to displaying environmentally regulated frameshifts are enriched in non-optimal codons.

**Figure 6.**
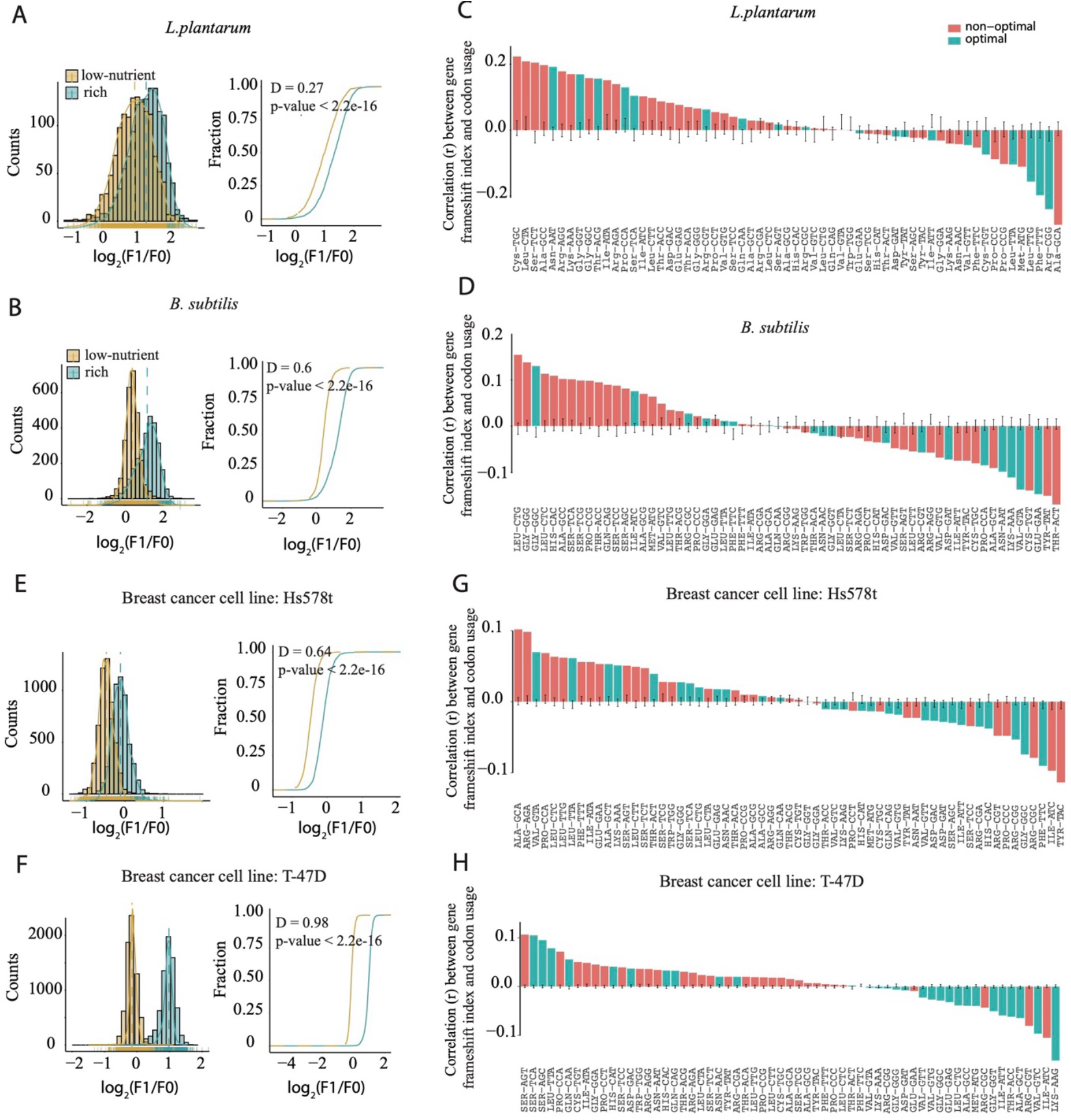
Out-of-frame co-translational decay of mRNA induced in bacteria upon starvation. (A) Distribution of gene-specific frameshift index for *L. plantarum* growing in rich (MRS broth) and after transferring it to low-nutrient conditions (0.5xLB media) for 15 mins (Data obtained from Huch, *et.al*^9^). A cumulative fraction plot with the frameshift index distribution is showed on the right. Statistical analysis is performed using Kolmogorov–Smirnov test with P-value and D as distance between control and treatment distributions. (B) as A, but for *B. subtilis* exponentially growing in rich media (LB) and in low-nutrient conditions (minimal growth medium) (see in Methods section). (C) Pearson correlation between gene-specific frameshift index and codon usage. Only genes (log_2_(F1/F0_) control_ > 0 and log_2_(F1/F0) _treatment_ < 0) in *L. plantarum* were used for calculating correlations. Codons defined as optimal (in green) or non-optimal (in red) codons according to codon adaptation index, obtained from Fuglsang^51^. (D) as C, but for *B. subtilis*. tRNA adaptation index were obtained from Perach *et.al*^50^. (E) as A, but for cell line Hs578t growing in rich media (DMEM medium) and in glutamine deprived conditions. (Data obtained from Loayza-Puch, *et.al*^36^). (F) as A, but for cell line T-47D. (G) as C, but for Hs578t. Optimal codons and non-optimal codons are shown in green and red, respectively according to Forrest *et.al* ^61^. (H) as G, but for cell line T-47D.

Lastly, we expanded our analysis to explore whether accelerated co-translational mRNA decay via ribosome frameshifting in response to low nutrient conditions also occurs in human cells (Fig 6E-H). We reanalysed ribosome profiling data from basal and luminal breast cancer cell lines after 48h growth in glutamine-free medium^36^. All analysed cell lines exhibited a significant increase in *out-of-frame* ribosome protection patterns (Fig S6D). This was particularly clear for Hs578t (frameshift transcripts increased from 3237 to 5222 transcripts, from 71% to 99% of detected transcripts) and T-47D (frameshift transcripts increased from 13 to 5830, from 0.1% to 83% of detected transcripts) (Fig 6E-F, Fig S6E). Overall, our findings show that environmentally induced ribosome frameshifting and out-of-frame mRNA decay is a conserved process from bacteria to humans.

### NMD limits cellular growth in low nutrient condition

Having observed that accelerated global mRNA decay via ribosome frameshifting in response to low nutrient conditions is general in biology, we wondered in which conditions this would be advantageous for the cells. Our data suggest that cells can repurpose the NMD degradation machinery to limit mRNA abundance. We hypothesize that this should also restrict translation and thus cell growth. To test this, we first compared the maximum growth rate of exponentially growing cells with (wild type) and without active NMD (*upf1Δ*) (Fig S3C). Despite their similar growth rates, we observed a marginal increase in growth rate for the *upf1Δ* strain compared to the wild type in CSM conditions. Additionally, wild-type cells exhibited an extended ‘lag phase’ (the phase without replication) relative to the *upf1Δ* strain, particularly under low nutrient conditions (Fig 7A). These findings suggest that under suboptimal growth conditions, NMD plays a crucial role in restraining cell growth.

**Figure 7.**
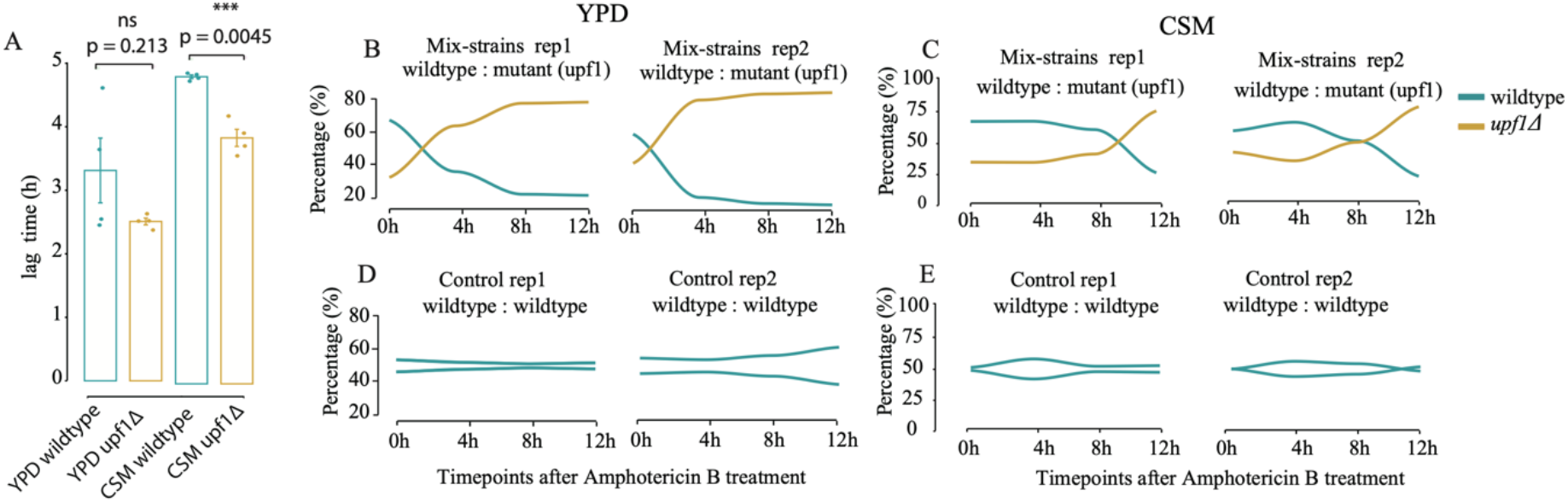
NMD limits cell proliferation in low nutrient condition. **(A)** Bar plot comparing the lag time (h) between wildtype (BY4741) and NMD mutant (*upf1Δ*) both in YPD and CSM. Each condition consists of four biological replicates. Statistical analysis was performed using two-sided t tests. (B) The proportions dynamics of wildtype (BY4741) and NMD mutant (*upf1Δ*) strains in co-culture following a three-hour exposure to amphotericin B (10 mg/l) and subsequent transfer to YPD. The proportions were recorded at four-hour intervals for a total duration of 12 hours post-treatment. (C) as B, but for co-culture in CSM with a subsequent transfer to CSM. (D) as B, but for co-culture wildtype (BY4741) and another wildtype (BY4741) with different barcodes as the control. (E) as C, but for the control in CSM.

Restricting cell growth and conserving limiting resources during nutrient deprivation is reminiscent of the bacterial *stringent response* that allows cells to survive the metabolic stress and enter a dormancy state in a reversible way^37^. Thus, we speculated whether NMD might also facilitate the transition of eukaryotic cells into a quiescent state under nutrient scarcity. To explore this possibility, we evaluated how wildtype cells (active NMD) and cells deficient in NMD (*upf1Δ*) recover and resume growth in the respective media following exposure to antifungal agents with a lethal dose of a drug (killing >99% of cells). We conducted an experiment where co-cultures of genetically tagged wildtype and *upf1Δ* cells (mixed in a 1:1 ratio) were treated with amphotericin B (10 mg/l) for three hours. Subsequently, these cells were transferred to fresh media (YPD and CSM, respectively), and we tracked the shifts in the proportions of each strain at four-hour intervals, as shown in Fig7B-E. Even if all cultures remained in lag phase as measured by OD_600_ for > 12 h, we observed that the ratio of wildtype to *upf1Δ* cells in YPD started to shift under four hours (Fig 7B). However, in CSM, this alteration persisted for a longer duration, extending up to eight hours (Fig 7C). This trend was not observed in co-cultures of wildtype strains, regardless of YPD or CSM media (Fig 7D-E). We hypothesized that the observed phenotype might be an adaptive strategy where wild type cells conserve energy and resources by slowing down cell division in unfavourable environments.

## Discussion

Quality control pathways are essential to ensure that aberrant mRNAs or proteins are cleared out from the cells. In general, quality control mechanisms are energetically expensive, thus checkpoint mechanisms are in place to ensure that they only act on aberrant molecules. Here we have shown that mRNA quality control is repurposed to globally control mRNA stability during poor nutrient conditions. Our work shows that ribosomes induce massive genome-wide frameshift events by sensing limiting nutritional conditions. We hypothesize that his could be a new mechanism for ribosomes to easily control mRNA life by facilitating its decay. Here, we have examined the position of ribosomes associated with mRNA decay and identified that about 77% of *S. cerevisiae* transcriptome is undergoing -1 frameshift-associated decay under low nutrition conditions. Although this phenomenon is obvious in poor media, a small fraction of genes also show preferential *out-of-frame* 5’P degradation signatures in optimal growth conditions. This suggests that *out-of-frame* decay plays a central role in gene expression and that, despite the known ability of NMD to target canonical mRNAs^14,15,16^, the magnitude of its effect has likely been underestimated. We suspect that it is due to the fact that previous work focused mainly on the bulk of translating ribosomes, and ignored the subset of mRNAs undergoing degradation that are transient and difficult to study. Importantly, this environmentally induced *out-of-frame* decay was also evident when investigating the mRNA turnover using RNA metabolic labelling. Using SLAM-Seq^32^ we estimated that NMD associated decay accounts for a minimum of 32% of the total decay rate in low-nutrition conditions.

To understand how ribosomes induce genome-wide -1 frameshifts in poor nutritional conditions, we investigated the potential involvement of known programmed ribosome frameshift (PRF) sites. –1 PRF are often associated with the presence of an heptanucleotide slippery sequence (X XXY YYZ)^38^ and a downstream secondary structure element^39^. The slippery sequence promotes ribosome slippage into the alternative frame, enabling codon-anticodon interactions in both frames, while the secondary structure element serves to slow down the ribosome at that position. The more optimistic estimates suggest that PRF sites exist in around 10% of all protein-coding genes across organisms^21,22^. However, our work indicates that in our experimental conditions, the bulk of the transcriptome experiences a frameshift independent of known PRF sites (Fig S4, Table S4).

On the contrary, we highlight the role of low codon optimality in the appearance of environmentally induced ribosome frameshifts. This suggest that the molecular mechanism underlying the massive frameshifts that we report here could be related to the “hungry codon” frameshifts that occur during ribosome translocation via tRNA slippage of the P-site while the A site is vacant^19,20,40^. This frameshift mechanism can be coupled^41^ or be independent^40^ of *cis* elements enhancing its efficiency (*i.e*., secondary structure element) and is conserved from bacteria to mammals. Recent work has demonstrated that lower concentrations of charged tRNA^Gln-CUG^ enhances frameshifts associated with the CAG-encoded polyglutamine repeats in huntingtin (Htt)^42^. Similarly, it has been shown in mouse embryonic fibroblasts that amino acid deprivation can lead to selective uncharging of glutamine-specific tRNAs and increase of frameshifts in proteins containing polyglutamine tracks^43^. While previous studies focus on particular genes or tRNAs, our work demonstrates how a similar phenomenon can rewire gene expression by affecting the bulk of the transcriptome.

The model for environmentally induced frameshifts that we propose here suggests that the action of NMD will be key to regulate mRNA stability in those conditions (Fig 1D). Interestingly, NMD has been previously shown to target preferentially transcripts with lower codon optimality^30^. In fact, Celik *et al*. showed that NMD substrates tend to have a significantly higher ratio of *out-of-frame* reads as measured by ribosome profiling, something that further supports our working model. Here we show that NMD targeting through ribosome frameshifting is modulated by nutritional conditions. Furthermore, we show that the correlation between low codon optimality and frameshift associated decay is maintained in bacteria that lack canonical NMD^44^. This suggests that not only NMD, but general mRNA degradation and quality control mechanisms are used to transfer environmental information into mRNA decay signatures via ribosome frameshifting in both eukaryotes and prokaryotes. Interestingly the mechanism that we report here is different from the previously reported role of ribosome collisions in signal transduction^45^. In fact, in our conditions we do not observe an increase of disomes but rather a decrease (Fig S1C). We hypothesize that, when low nutrition conditions are maintained over time, both translation initiation and elongation are limited. This would reduce the likelihood for ribosome collisions, but not affect the frameshift mediated mechanism that we propose here.

In addition to mRNA degradation, the massive frameshifting we describe can be expected to affect the proteome. Decreasing the abundance of available mRNA templates for translation will directly affect protein synthesis. As expected, we observe that genes with a higher level of frameshifts present a lower relative protein abundance than those with lower frameshifts (Fig 5C). Our observations are congruent with previous work investigating alternative mRNA quality control pathways. For example, it has previously been demonstrated that ribosome stalling at non-optimal codons can reduce protein synthesis rates by increasing mRNA decay rates via No-Go Decay^11^. As we observe a decrease of ribosome collisions during continued growth in low nutrients (Fig S1C), our work suggests that frameshifts and not only ribosome collisions can transduce nutritional information into gene expression regulation. The regulation of protein abundance via frameshifting and increased mRNA decay further supports the role of the ribosomes as nutritional sensors for dynamic regulation of a functional and balanced proteome. Our data showing that NMD can restrict cellular growth in low nutrient conditions suggest that in eukaryotes, nutritional sensing by the ribosomes followed by NMD decay could exert a function similar to the stringent response in bacteria. Namely, allowing cells to conserve limiting resources during nutrient deprivation, survive the metabolic stress and enter a dormancy state in a reversible way^37^.

Altogether, our work suggests that the translation process can act as a sensor for the dynamic regulation of a functional and balanced proteome by directly regulating mRNA life. Increased mRNA decay under poor nutritional conditions will limit energy used for translation. This raises the possibility that frameshifting may be beneficial for cells also in terms of releasing ribosomes and facilitating the degradation of peptides for subsequent recycling in response to low nutrition conditions. Finally, faster mRNA turnover should also facilitate rewiring of the transcriptome and a swifter adaptation to new environments, something especially beneficial for unicellular organisms.

## Limitations of the study

We have shown that environmentally induced frameshift is general in biology. In addition to the general response regarding mRNA stability, frameshift events during translation elongation can be expected to generate aberrant proteins with canonical N-terminal polypeptide sequence and altered C-terminal regions. Although those events are expected to be very rare, the presence of proteins with altered C-terminal regions could lead to the increase of protein aggregates or other proteostasis problems. However, further work will be needed to determine the potential impact on cell physiology of the putative generated frameshifted peptides. Our work also proposes a model where NMD limits cell growth in low nutrition conditions. This raises the appealing possibility that ribosome frameshifting followed by NMD decay could facilitate the entry into quiescence for eukaryotic cells. We hypothesize that in eukaryotes NMD could exert an analogous function to the type II RelE/RelB toxin-antitoxin system in bacteria that leads to a generalized mRNA degradation in response to low nutrition conditions^46,47^. This new mechanism could be relevant to understand how eukaryotic cells could escape antifungal treatment or even chemotherapy. However, additional work would be required to explore this hypothesis.

## ACKNOWLEDGEMENTS

We thank all members of the Pelechano, Kutter and Friedländer laboratories for useful discussions and suggestions. We thank, José E. Pérez Ortín for additional feedback. Computational analysis was performed on resources provided by SNIC through Uppsala Multidisciplinary Center for Advanced Computational Science. This project was funded by the Swedish Foundation’s Starting Grant (Ragnar Söderberg Foundation), a Wallenberg Academy Fellowship [2016.0123 and 2021.0167], the Swedish Research Council [VR 2016-01842, 2020-01480 and 2019-02335], Karolinska Institutet (SciLifeLab Fellowship, SFO and KI funds) a Joint China-Sweden mobility grant (STINT, CH2018-7750) and a Humboldt Research Fellowship for Experienced Researchers (SWE 1221518 HFST-E) to V.P. Y.Z. was funded by a fellowship from the China Scholarship Council. LN acknowledges funding from the EU H2020-MSCA-IF-2018 program under grant agreement [845495 - TERMINATOR] and SC RA grant agreement [21SCG-1F006]. IP receives funding from the Helmholtz Young Investigators program of the Helmholtz Association and from the European Research Council (ERC) under the European Union’s Horizon 2020 research and innovation program (grant agreement ERC-STG No 948544). PS is the recipient of grants from the Swedish Cancer Fund (19-0133 and 22-2014).

## AUTHOR CONTRIBUTIONS

VP, YZ and LN conceived the project. YZ performed most experimental and computational work. LN contributed to the frameshift analysis and discussion. EF performed the proteomic analysis under the supervision of IP. CB contributed to proteomic data analysis. IA provided additional experimental assistance. SH contributed to polysome isolation and discussion. EG and PS contributed to initial experiments and discussion. YZ and VP drafted the original manuscript. All authors reviewed and edited the manuscript. VP supervised the project.

## Declaration of Interests

VP, SH and LN are co-founders and shareholders of 3N Bio AB which has filed a patent application regarding the study of the microbial degradome. All other authors declare no competing interests.

## Methods

### Strains and growth conditions

All yeast experiments were performed using *Saccharomyces cerevisiae* strain BY4741 (MATa *his3Δ1 leu2Δ0 met15Δ0 ura3Δ0*) if not stated otherwise. *S. cerevisiae* was grown to mid exponential phase (OD_600_∼0.6) at 30 °C with rotati using YPD (1% yeast extract, 2% peptone, 2% glucose), Complete Supplement Mixture (CSM from Formedium™) or Synthetic Complete (SC from MP Biomedicals™) medium (composition details seen in Table S1).

To measure the growth curve, cells were initially pre-cultivated at 30°C with rotation overnight until reaching the mid-exponential phase (OD_600_∼0.6). Subsequently, the cells were diluted to an OD_600_ of approximately 0.01 in preparation for measurement using a plate reader. OD_600_ was measured every 10 minutes with 5 minutes of pre-shaking before each measurement. This process continued for a total duration of 36 hours.

*Bacillus subtilis* 168 (trpC2) strain was pre-cultivated at 37°C with rotation in LB and minimal growth medium (per 50 ml: 5X minimal salts solution,10 ml; glucose (50% (w/v)), 0.5 ml; casamino acids (2% (w/v)), 0.5 ml; tryptophan (10 mg/ml), 0.1 ml; iron ammonium citrate (2.2 mg/ml), 0.05 ml, deionized water, 39 ml) with tryptophan supplementation at OD600 < 0.8. Cultures were diluted to OD_600_∼0.05 and collected until reaching the mid-exponential phase (OD_600_∼0.6).

For stress treatments in *S. cerevisiae*, we grew the cells to exponential phase, then split them into different flasks for different stress treatments. For heat shock, cells were exposed to 42°C for 30 minutes. For oxidative stress, cells were treated with 0.2 mM H_2_O_2_ for either 5 or 30 minutes. To induce amino acid deprivation, cells were quickly collected by centrifugation and then washed with CSM medium lacking amino acids. The cells were then incubated for 30 minutes (starting from the time they were first exposed to the amino acid deprived CSM medium). For glucose deprivation, mid exponential *S. cerevisiae* cultures at OD_600_ ∼0.8 were spun down, washed twice, and resuspended in pre-warmed YP (lacking glucose) and then grown at 30 °C for 5 minutes and 15 minutes as time points. To induce ribosome collisions in *S. cerevisiae*, we grew the cells to exponential phase in SC medium and then transfer them to SC media without histidine medium for 30 minutes. For addition of single amino acid experiment, arginine, proline, or serine were individually added as single amino acids to reach the comparable concentration as present in SC medium. Specifically, we increased the concentration of each amino acid from 50 to 85.6 mg/L for arginine, and from 0 to 85.6 mg/L in the case of proline and serine. *S. cerevisiae* was inoculated into each medium overnight as precultures. Cultures were diluted to OD600∼0.05 and collected until reaching the mid-exponential phase (OD_600_∼0.6) at 30°C.

All yeast and bacteria samples prepared for HT5Pseq libraries were collected through centrifugation and preserved by freezing them in liquid nitrogen. Total RNA was isolated by the standard phenol: chloroform method and DNA was removed by DNase I treatment. RNA concentration was measured with Qubit and RNA quality was checked by 1.2% agarose gel or on a BioAnalyzer using an RNA Nano 6000 chip (Agilent Technologies).

### HT-5Pseq library preparation

HT-5Pseq libraries were prepared following a previously established protocol^25^ if not stated otherwise. Briefly, 6 μg of total RNA was subjected to DNase treatment, and the resulting DNA-free total RNA was ligated with an RNA oligo containing UMI (rP5_RND oligo). The ligated RNA was reverse transcribed using Illumina PE2 compatible oligos with random hexamers and oligo-dT as primers. To eliminate RNA in RNA/DNA hybrids, samples were treated with sodium hydroxide at 65°C for 20 minutes. Ribosomal RNAs were depleted using DSN (Duplex-specific nuclease) and a mixture of ribosomal DNA probes. To deplete Ribosomal RNAs in *B. subtilis* 168 (trpC2), we used the in-house rRNA DNA oligo depletion mixes described previously^9^. Finally, the samples were amplified by PCR and sequenced on an Illumina NextSeq 2000 instrument using an average of 45 cycles for read1.

### HT5Pseq data processing and analysis

HT5Pseq reads were demultiplexed using bcl2fastq (v2) and the 3’’-sequencing adapter was trimmed using cutadapt V4.0. After that, the 8-nt random barcodes located on the 5’ ends of reads were extracted and added to the fastq file header as UMIs using UMI-tools^48^. Reads were mapped to the reference genome (SGD R64-1-1 for *S. cerevisiae* and GCF_000009045.1_ASM904v1 for *B. subtilis*) using star/2.7.0^49^ with the parameter --alignEndsType Extend5pOfRead1 to exclude soft-clipped bases on the 5’ end. Duplicated 5’ ends of read introduced by PCR during library preparation were removed based on random barcodes sequences using UMI-tools. Analysis of 5’ end positions was performed using the *fivepseq* package^24^ version 1.2.0. This included analysis of the relative positions of the 5’ ends of the mRNA reads with respect to the start codon, stop codon, and codon-specific pausing. Specifically, the unique 5’mRNA reads in biological samples were averaged and normalized to reads per million (rpm). The relative position of 5’ mRNA reads at each codon were summed up and used for all additional analyses. Metagene plots for frame (F0, F1 or F2) preference are shown as the average sum (in rpm) of each frame over a sliding window of 20 codons, for +/- 100 bp from the start and end, and for +/- 300 bp from the middle of CDS for each gene. *Fivepseq* output files will be deposited at the SciLifeLab Data repository.

To calculate gene-specific frameshift index, we summed the reads for each frame of every transcript after excluding the first and last 50 nt. Transcripts with greater than 20 reads per million (rpm) were considered for subsequent analysis in each biological replicate. The frameshift index of each transcript was determined by dividing the number of in-frame reads (F1) by out-of-frame reads (F0), with the result expressed in logarithmic scale. Only transcripts that were present in all biological replicates were included in the calculation of frameshift index. Similarly, the frameshift index for each codon was calculated by dividing the in-frame reads at position -17 (F1) by the out-of-frame reads at position -18 (F0), with the result expressed in logarithmic scale. High frameshift genes are defined as: log_2_(F1/F0) _YPD_ > 0.2 and log_2_(F1/F0) _CSM_ < -0.2) (according to mean value in each distribution) and non-frameshift genes are defined as: log_2_(F1/F0) _YPD, CSM_ > 0).

Disomes were detected by the *fivepseq* package ^24^ in the queue statistics output using default parameters. For each gene, *fivepseq* first determines windows of periodicity 30 nt +/-3.6 nt that have a Fast Fourier transform (FFT) signal of more than 20. The identified windows are then continuously extended and/or merged to bigger windows until the periodicity signal is no longer increasing. The merged windows are then sliced to a range between the two tallest peaks. Those peaks are determined based on the p value of a count falling withing a Poisson distribution with a lambda corresponding to the average count in the given range. Counts with a p value less than 0.001 are considered peaks. In the context of FFT signals, a periodicity of 30 nt ± 3.6 nt indicates the occurrence of two collided ribosomes (disomes). This periodicity indicates the presence of 2 adjacent protection sites separated exactly 30 nt (that is the distance covered by a full ribosome). Similarly, a length of 60 nt ± 3.6 nt corresponds to the presence of three ribosomes, and this pattern continues for subsequent lengths.

To determine the proportion of the transcriptome degraded through *out-of-frame* co-translational decay in CSM, we used a simulation in which we mixed the codon protection index^8^, defined as log_2_(F1/((F2+F0)/2)) from two samples at different ratios. Specifically, we combined the in-frame decay data from YPD (assuming a 100% of in-frame degradation) with simulated *out-of-frame* decay data (by shifting the YPD data by -1nt to generate a theoretical 100% *out-of-frame* decay). The use of the codon protection index instead of the frameshift index in this context is based on the fact that the YPD data shifted by -1nt, resulting in a change in all frames to the new corresponding -1nt frames. By using the codon protection index, we ensure that all frame changes are taken into consideration and properly accounted for in the analysis. We mixed both samples at different ratios (increasing 10% *out-of-frame* decay at each mixing) to estimate the median of each codon protection index distribution. Each mixing process was iterated 1000 times to obtain 95% confidence intervals. Finally, we used the generated distribution of codon protection indexes to estimate the percentage of *out-of-frame* co-translational decay. To analyse whether the frameshift is induced by the ribosome programmed frameshift slippery motif, we utilized putative PRF sites downloaded from PRFdb^22^ (Supplementary Data S4). 5PSeq reads were aligned to the annotated slippery sequences motifs with an extension of 99 bp both upstream and downstream. We calculated the coverage for each frame by applying a sliding window of 3 nucleotides and taking the average. Finally, we plotted the calculated proportion for each frame.

To analyse frameshift in *B. subtilis* 168 trpC2 at early stationary phase and *L. plantarum* in low nutrients, we obtained the dataset from Huch *et.al* ^9^ with GEO accession number: GSE153497. The calculation of frameshift index for each gene was performed as described above, with the only modifications of using *in-frame* reads at position -14 (F1) and the *out-of-frame* reads at position -15 (F0) due to the difference of ribosomal fragment protection size between yeast and bacteria. Transcripts with greater than 30 reads and 10 reads per million (rpm) were considered for subsequent analysis in each biological replicate.

Gene-specific frameshift index for *B. subtilis* 168 and *L. plantarum* were computed as follows: log2(F1/F0)_control_ > 0 and log2(F1/F0) _treatment_ < 0. Statistical analysis for frameshift index distribution between two populations was performed with a Kolmogorov–Smirnov test.

To infer codon optimality in *B. subtilis* 168 *and L. plantarum*, we obtained tRNA adaptation index and Codon adaptation index from Perach *et.al* ^50^ and Fuglsang^51^ respectively.

All clustering analyses and heatmap were performed by k-means using Complexheatmap packages from R and Bioconductor^52^. Gene Ontology enrichment analysis were performed with ClusterProfiler^53^ using Fisher’s exact test with p adjusted value < 0.05. Datasets for *S. cerevisiae* tRNA adaptation index, mRNA codon stability index, translation efficiency were obtained from Carneiro *et al*.^54^, gene expression level, mRNA half-life, 3’/5’ UTR length were obtained from Xu *et al*. ^55^, Presnyak *et al*.^10^ and Pelechano, *et.al*^56^ respectively.

### SLAM-Seq metabolic labeling

Metabolic labelling of newly synthesized RNA molecules was performed as previously described^32,57,58^. Briefly, 4-Thiouracil (4tU, Sigma) was dissolved in NaOH (83 mM) used for labelling RNA molecules. The final concentration of 4tU for YPD and CSM was 5 mM^57^ and 0.2 mM^58^, respectively. MES buffer (pH 5.9) with a final concentration of 10 mM was prepared with media to avoid the pH change due to 4tU addition. Wildtype (BY4741) and NMD mutant (*upf1*Δ) were used for these experiments. To perform pulse and chase experiment, RNA molecules were labelled with 4tU (prepared as above) for 1h during cell grown at mid exponential phase. Cells were washed and resuspended in prewarmed medium (YPD with MES buffer or CSM with MES buffer) without 4tU and time points were collected at 0, 15 and 30 mins by centrifugation and snap frozen in liquid nitrogen immediately.

To perform thiol(SH)-linked alkylation, the reaction including 5 μg of RNA was incubated with iodoacetamide (final concentration at 0.5 M) at 50°C for 15 minutes^32^. The reaction was stopped by adding 0.1 M DTT (final concentration at 20 mM). RNA was then precipitated by 3 M of sodium acetate and pure ethanol. Purified RNA was subjected for ribosomal RNA depletion using RiboPools Depletion Kit (siTOOLs Biotech). Libraries were prepared by Ultra™ II Directional RNA Library Prep Kit for Illumina® following manufacturer’s instructions. Sequencing was performed on Illumina Nextseq 2000 sequencer with single end read length for 150 cycles.

### SLAMseq data analysis

SLAM-Seq data analysed was performed with slamdunk (v0.4.3) provided by nfcore pipeline(v1.0.0)(https://nf-co.re/slamseq). Firstly, fastq compressed files (fastq.gz) were converted to reverse complementary reads method and feed into slamdunk nf core pipeline as stranded libraries were prepared with dUTP. Adapter contamination and low-quality bases were trimmed using TrimGalore(v 0.6.4) (trim length 27bp). Slamdunk was used for mapping, quantifying nucleotide-conversions and collapsing quantifications on gene level. At least 1 T>C conversions per read was regarded as a confident call for labelled RNA reads. Genes with less than 20 reads were filter out. SNP masking was employed to distinguish filter T>C SNPs from converted nucleotides. T > C reads counts in each library were normalized to total library read counts. Normalized T > C reads counts across time were fitted to non-linear decay model equations with R function nls to calculate degradation rate for each condition: y ∼ C*exp(−a*t_m_) where normalized T > C reads counts fitted to y, with chasing time points from 0, 15 and 30 mins being fitted to t_m_. The model was used to estimate parameters C and a, from which the RNA half-life was calculated as log2(a). Degradation rate was calculated as 60 * ln(2) / t_1/2_ (t_1/2_ is half-time of gene). To show the percentage of labelled read across time, T>C conversions were normalized to chase-onset (t_0_). Environmental NMD-sensitive genes are defined according to degradation rate in wildtype and mutant: degradation rate (wt/upf1*Δ*)_CSM_ > 1.2 and (wt/ upf1*Δ*)_YPD_ < 0.8).

### Ribosome profiling library preparation

Yeast overnight cultures were diluted to OD_600_ 0.05 in 1000 ml CSM medium and grown at 30 °C. All cells were collected by vacuum filtration and freezing by liquid nitrogen. To perform lysis, 200 µl glass beads and 500 µl of freshly prepared lysis buffer (composed of 20 mM Tris pH 8.0, 140 mM KCl, 1.5 mM MgCl_2_, 1% Triton X-100, 0.1 mg/mL cycloheximide, 2 mM DTT, and an EDTA-free protease inhibitor tablet) were added. Cells were pulverized by vortexing using a Multimixer for 2 minutes, followed by a 5 minute incubation on ice. This process was repeated three times to maximize the yield of RNA. After adding an additional 200 µl of lysis buffer to compensate for volume loss, the lysate was then centrifuged at 3000 g for 5 minutes at 4°C. After quantifying RNA concentration, 200 µg RNA was incubated with 2 µl 100 U/µl RNase I (Ambion) for 1 h at 22 °C with gentle agitation (700 rpm) and the reaction was inhibited by the addition of 15 µl SuperaseIn (Thermo Fisher). Another 50 µg RNA from the lysate was kept on ice undigested to use for polysome profiles. Linear sucrose gradient centrifugation and fractionation were then performed as described previously^59^, except we modified the 10X sucrose gradient buffer (consisting of 0.5 M Tris-Acetate pH 7.0, 0.5 M NH_4_Cl, and 0.12 M MgCl_2_) to make it suitable for yeast cells. RNA was extracted in 20 mM Tris-HCl (pH 7.5), 300 mM sodium acetate, 2 mM EDTA, 0.25% SDS overnight at room temperature with rotating and then precipitated. Monosome fraction RNA was isolated by the standard phenol: chloroform method. RNA footprints were isolated at 20-35 nucleotide size using 15% TBE-urea gel.

To further ligate 3’ and 5’ adapters, RNA samples were treated with PNK to obtain 5’ phosphorylated and 3’ hydroxylated ends, followed by ribosomal RNA depletion by using the RiboPools Depletion Kit (siTOOLs Biotech). For total RNA, ribosomal RNA was depleted (as described above) followed by fragmentation with magnesium at 80°C for 5 mins. RNA samples were treated with PNK to obtain 3’ hydroxylated ends for further 3’ ligation. Monosome RNA footprints and total RNA were prepared using a Small RNA Sample Preparation Kit for NEXTFLEX^®^ following manufacture instructions. Sequencing was performed with single end setting, read length 47 bp on Illumina Nextseq 2000 sequencer.

### Ribosome profiling data analysis

Ribosome profiling data was trimmed using cutadapt and the following parameters: -a TGGAATTCTCGGGTGCCAAGG. To keep the minimum length, the cutoff set to 10 nt. After that the 4-nt random barcodes from both 5’ and 3’ ends of reads were extracted as UMIs. Ribo-seq reads and RNA seq reads were selected at 28-32 nt and 20-70 nt, respectively. Reads were mapped to reference genome (SGD R64-1-1) using star/2.7.0. Analysis of 5’ end positions was performed using *Fivepseq* package as described above. The distribution of frames with respect to the size of the ribosome-protected fragments were determined using the riboSeqR package in R (v4.0.4).

Ribosome profiling data for amino acid deprivation in human cell lines was obtained from Darnell, *et.al* with GEO accession number: GSE113751^60^. Ribosome profiling data were adapter trimmed using cutadapt by the following parameters: -a AAAAAAAAAA --minimum-length=13. Reads were mapped to reference genome (GRCh38) using star/2.7.0. Analysis of 5’ ends positions was performed using *Fivepseq* package as described above. The calculation of frameshift index for each gene were described above, with modifications as follows: 1) using in-frame reads at position -17 (F1) and the out-of-frame reads at position -18 (F0) according to human ribosome protection fragment size. 2) Only genes whose length was divisible by 3 and whose coding sequence (CDS) started with ATG and ended with either TAG, TGA, or TAA stop codons were considered. Codon optimality data for 293t was downloaded from Forrest, *et.al*.^61^

### Sample Preparation for proteomics analysis

Yeast cells grown in YPD or CSM media were quenched by adding pure trichloroacetic acid (Sigma Aldrich) to the yeast cultures to a final concentration of 10% (v/v) and incubating for 10 min on ice. Samples were then centrifuged at 2500 g for 5 min at 4°C and the supernatant was discarded. The pellet was washed twice with 10 ml cold acetone before being transferred into a new tube. After an additional centrifugation step at 3000 g for 5 min at 4°C, the acetone was removed and the pellet was further processed for protein extraction.

### Cell Lysis and Protein Extraction for proteomics analysis

To lyse the cells, cell pellets were first mixed with glass beads (Sigma Aldrich) and 500 μl of lysis buffer containing 8M urea, 50 mM ammonium bicarbonate and 5 mM EDTA (pH 8). The mixture was then transferred to a FastPrep-24TM 5G Instrument (MP Biomedicals) where cells were disrupted at 4°C by 4 rounds of bead-beating at 30 seconds with 120 seconds pause between the runs. Samples were then centrifuged for 10 min at 21’000 x g to remove cell debris and the supernatants were transferred into a new tube. The protein concentration was determined using the bicinchoninic acid Protein Assay Kit (Thermo Scientific) following the manufacturer’s protocol.

### In-solution protein digestion for proteomics analysis

100 µg of protein extracts were subjected to digestion. Samples were vortexed and sonicated for 5 min. In the first step, dithiothreitol (Sigma Aldrich) was added to a final concentration of 5 mM and incubated for 30 min at 37 °C to reduce the disulfide bridges followed by the alkylation of free cysteine residues with iodoacetamide (Sigma Aldrich) at 40 mM final concentration (30 min at 25°C in the dark). Samples were pre-digested with lysyl endopeptidase (Wako Chemicals) at an enzyme substrate ratio of 1:100 for 4 h at 37°C and then diluted 1:5 with freshly prepared 0.1 M ammonium bicarbonate to reduce urea concentration to 1.6M. Sequencing grade trypsin (Promega) was added at an enzyme substrate ratio of 1:100 and digested at 37°C for 16h. The digestion was stopped by adding formic acid (Sigma Aldrich) to a final concentration of 2%. The digested samples were loaded onto SepPak C18 columns (Waters) that were previously primed with 100% methanol, washed with 80% acetonitrile (ACN, Sigma Aldrich), 0.1% FA and equilibrated 3 times with a 1% ACN, 0.1% FA solution. The flow-through was loaded once more onto the columns and the peptides bound to C18 resins were afterwards washed 3 times with a 1% ACN, 0.1% FA solution and eluted twice with 300 μl 50% ACN, 0.1% FA. The elution was dried down in a vacuum centrifuge and peptides were resuspended in a 3% ACN, 0.1% FA solution to a concentration of 1 mg/ml before LC-MS analysis.

### Liquid chromatography–mass spectrometry (LC-MS) for proteomics analysis

Peptide samples were analysed in a Data-Independent Acquisition mode (DIA) with an Orbitrap Exploris 480 mass spectrometer (Thermo Fisher Scientific) equipped with a nano-electrospray ion source and a nano-flow LC system (Easy-nLC 1200, Thermo Fisher Scientific). Peptides were separated with a 50 cm fused silica capillary column with inner diameter of 75µm packed in house with 1.9 µm C18 beads (Dr. Maisch Reprosil-Pur 120). For LC fractionation, buffer A was 3% ACN and 0.1% FA and buffer B was 0.1% FA acid in 90% ACN and the peptides were separated by 2 h non-linear gradient at a flow rate of 250 nl/min with increasing volumes of buffer B mixed into buffer A. The DIA-MS acquisition method consisted of a survey MS1 scan from 350 to 1650 m/z at a resolution of 120,000 followed by the acquisition of DIA isolation windows. A total of 40 variable-width DIA segments were acquired at a resolution of 30,000. The DIA isolation setup included a 0.5 m/z overlap between windows.

### Quantitative proteomics data analysis

DIA-MS measurements were analysed with Spectronaut 16 (Biognosys AG) using direct searches. In brief, retention time prediction type was set to dynamic iRT (adapted variable iRT extraction width for varying iRT precision during the gradient) and correction factor for window 1. Mass calibration was set to local mass calibration. The false discovery rate (FDR) was set to 1% at both the peptide precursor and protein level. Digestion enzyme specificity was set to Trypsin/P and specific. Search criteria included carbamidomethylation of cysteine as a fixed modification, as well as oxidation of methionine and acetylation (protein N-terminus) as variable modifications. Up to 2 missed cleavages were allowed. The DIA-MS files were searched against the *Saccharomyces cerevisiae* proteome (UniProt version 2021-04-02). Differentially regulated proteins were determined with an unpaired t-test statistic with Storey method correction. After quantification of protein abundance, the proteins that were upregulated and downregulated were identified based on log_2_ (CSM/YPD) values >1 and < -1, respectively. Additionally, a Q-value of less than 0.01 was used as a criterion for selection. Volcano plot for protein abundance comparison was plotted using the EnhancedVolcano package from R. Gene set enrichment analysis (GSEA) for proteins was performed with ClusterProfiler^53^ using Fisher’s exact test with p adjusted value < 0.05. Protein-protein interactions were performed by metascape v3.5 with p adjusted value < 0.05.

## SUPPLEMENTARY FIGURES

**Figure S1.**
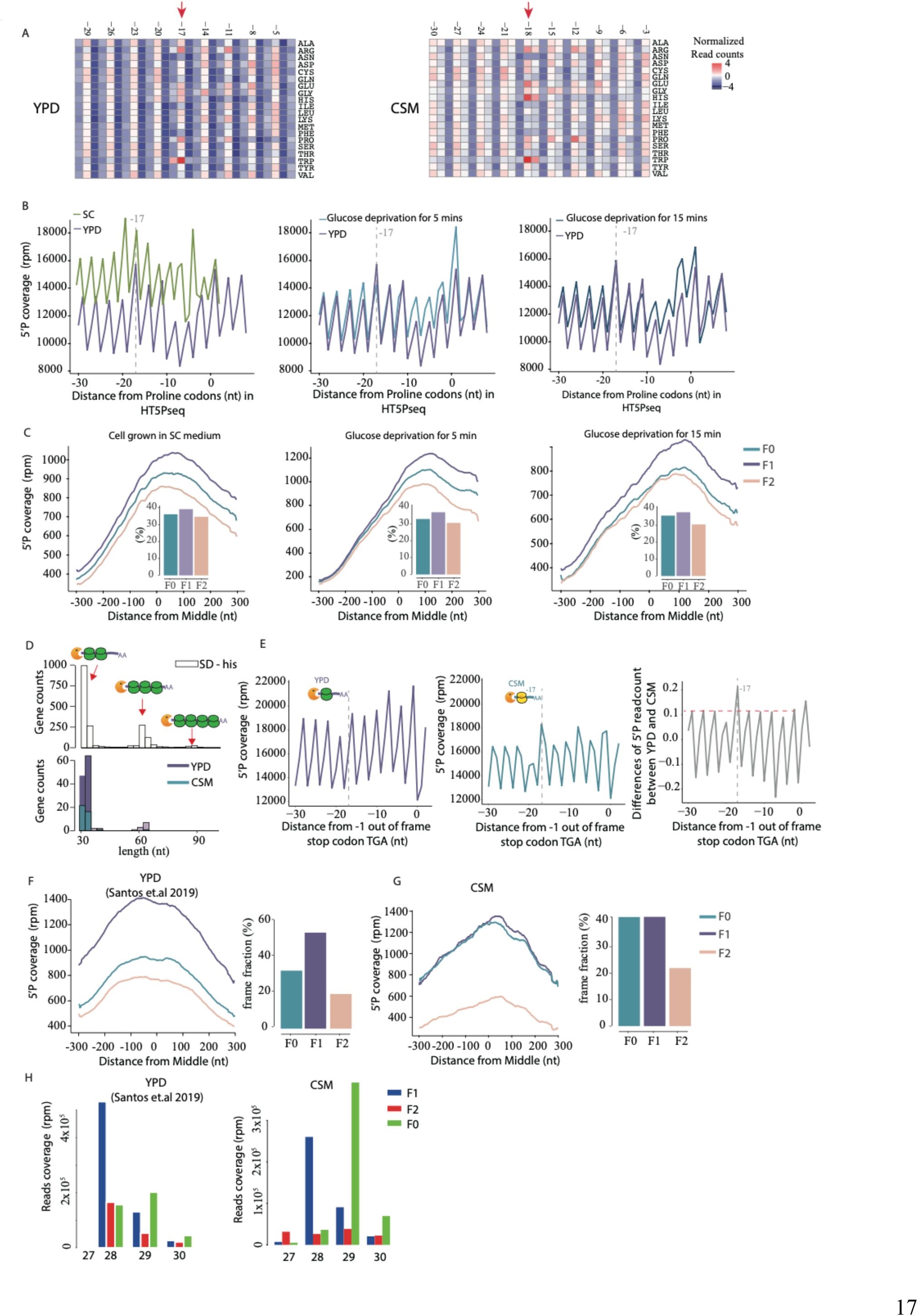
Co-translational mRNA decay reveals genome-wide -1 ribosome frameshift, Related to Figure 1. (A) Heatmap for metagene analysis displaying the 5′P reads coverage for each amino acids for YPD (left) and CSM (right). 5′P reads coverage is calculated by reads per million and the heatmap is row normalized. (B) Metagene analysis displaying the 5′P reads coverage for proline codons (CCG) in SC (in green, left), glucose deprivation for 5 mins (in blue, middle) and glucose deprivation for 15 mins (in dark blue, right) using HT5PSeq. 5′P reads coverage for proline codons in YPD (in purple) used as control in all conditions. Dotted lines at −17 corresponding to the *in-frame* 5′ end of protected ribosome located at the A site. (C) Relative 5′P coverage with average of 20 codons for each frame in SC (left), glucose deprivation for 5 mins (middle), glucose deprivation for 15 mins (right) from the middle of genes. The fractions calculated for each frame for metagene analysis (excluding the first and last 50 nt for each gene), are shown in the histogram. (D) Number of genes where ribosome collisions are detected by FFT analysis (see methods for details). A disome provides a signal at 30nt, a trisome at 60 nt and a tetrasome at 90nt. We show values for YPD, CSM and minimal media without histidine (SD-His). SD-His is shown as a positive control for inducing disomes^8^. Higher level of ribosome collisions are observed in YPD than in CSM growth. (E) Metagene analysis displaying the 5′P reads coverage for out-of-frame stop codons (*i.e*. TGA) in YPD (left), CSM (center) and normalized peak by dividing the relative peaks in CSM with in YPD (right) (See in Methods section). (F) as C, but for ribosome profiling in YPD. Data obtained from Santos *et.al* ^62^. (G) as C, but for ribosome profiling in CSM generated in this study. (H) The distributions (read counts in rpm) of frames in ribosome profiling for fragment sizes ranging from 27 to 30 nt are shown for YPD (left) and CSM (right, merged three replicates).

**Figure S2.**
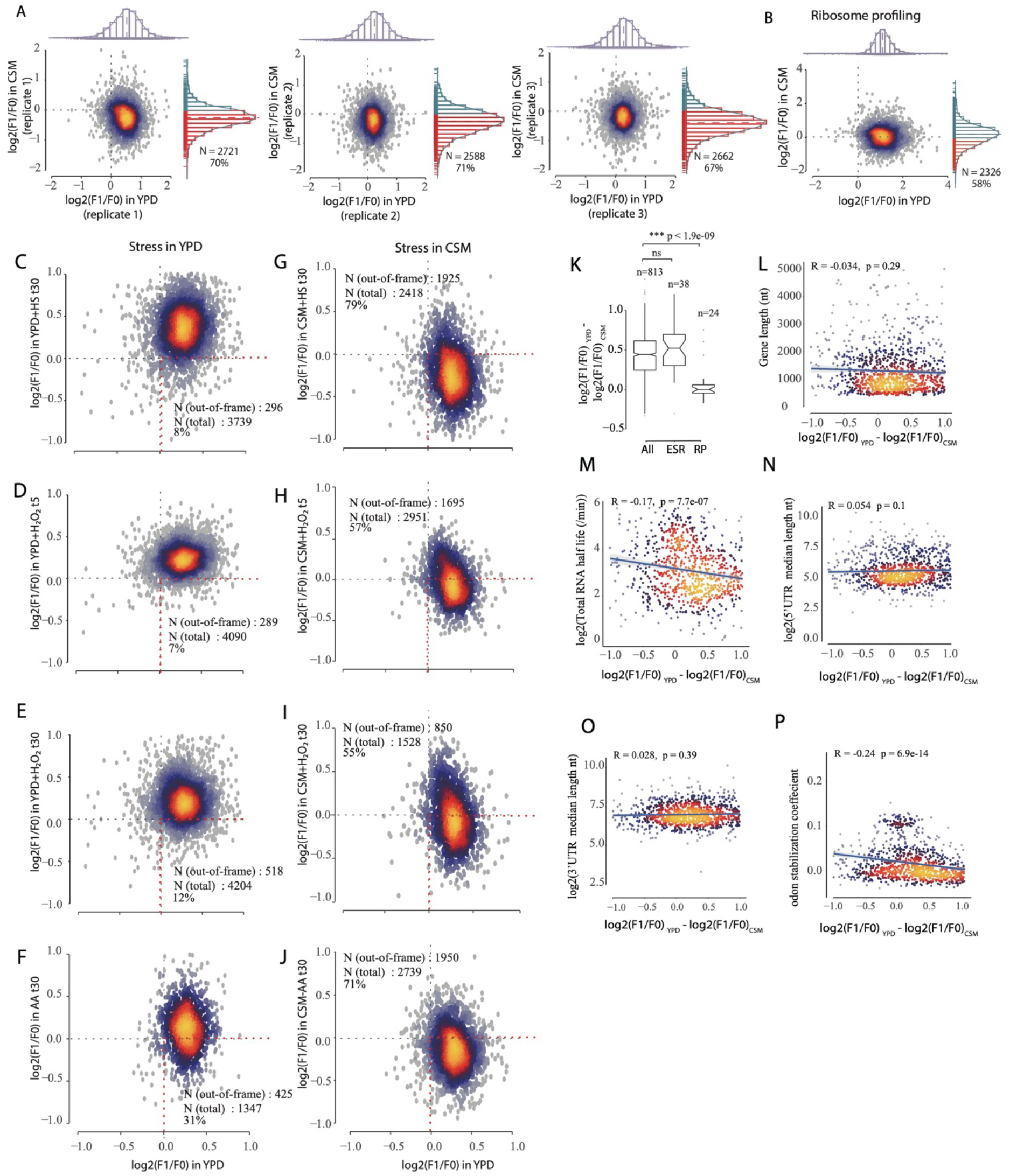
Genome-wide out-of-frame mRNA decay is environmentally regulated, Related to Figure 2. (A) Scatter plot comparing the frameshift index (log_2_(F1/F0)) of individual genes in YPD (x-axis) and CSM (y-axis) among three replicates. Histograms marked with red highlight genes that exhibited a mainly out-of-frame decay. Namely those genes with log_2_(F1/F0) < 0 in each replicate. (B) as (A), but for ribosome profiling. (C-F) as A, but for stress in YPD, *i.e*. (C) heat shock for 30 mins; (D) H_2_O_2_ (0.2 mM) exposure for 5 mins; (E) H_2_O_2_ (0.2 mM) exposure for 30 mins; (F) transfer from YPD to CSM medium lacking amino acids for 30 mins; (G-J) as C-F, but for cells grown long term in CSM. (K) Boxplot comparing frameshift index change for genes been detected among all stresses (All, N = 813), environmental response genes (ESR, N =38) and ribosomal protein genes (RP, N = 24) in Fig2C. Statistical analysis was performed using two-sided Wilcoxon rank-sum tests. (L-P) Spearman correlations between frameshift index change (x-axis) with RNA features, i.e. (L) gene length (nt); (M) Total RNA half-life in log scale (from Xu *et al* ^55^); (N) 5′UTR length (from Pelechano *et.al* ^56^); (O) 3′UTR length (from Pelechano *et.al* ^56^); (P) Codon stabilization coefficient (from Presnyak *et al* ^10^).

**Figure S3.**
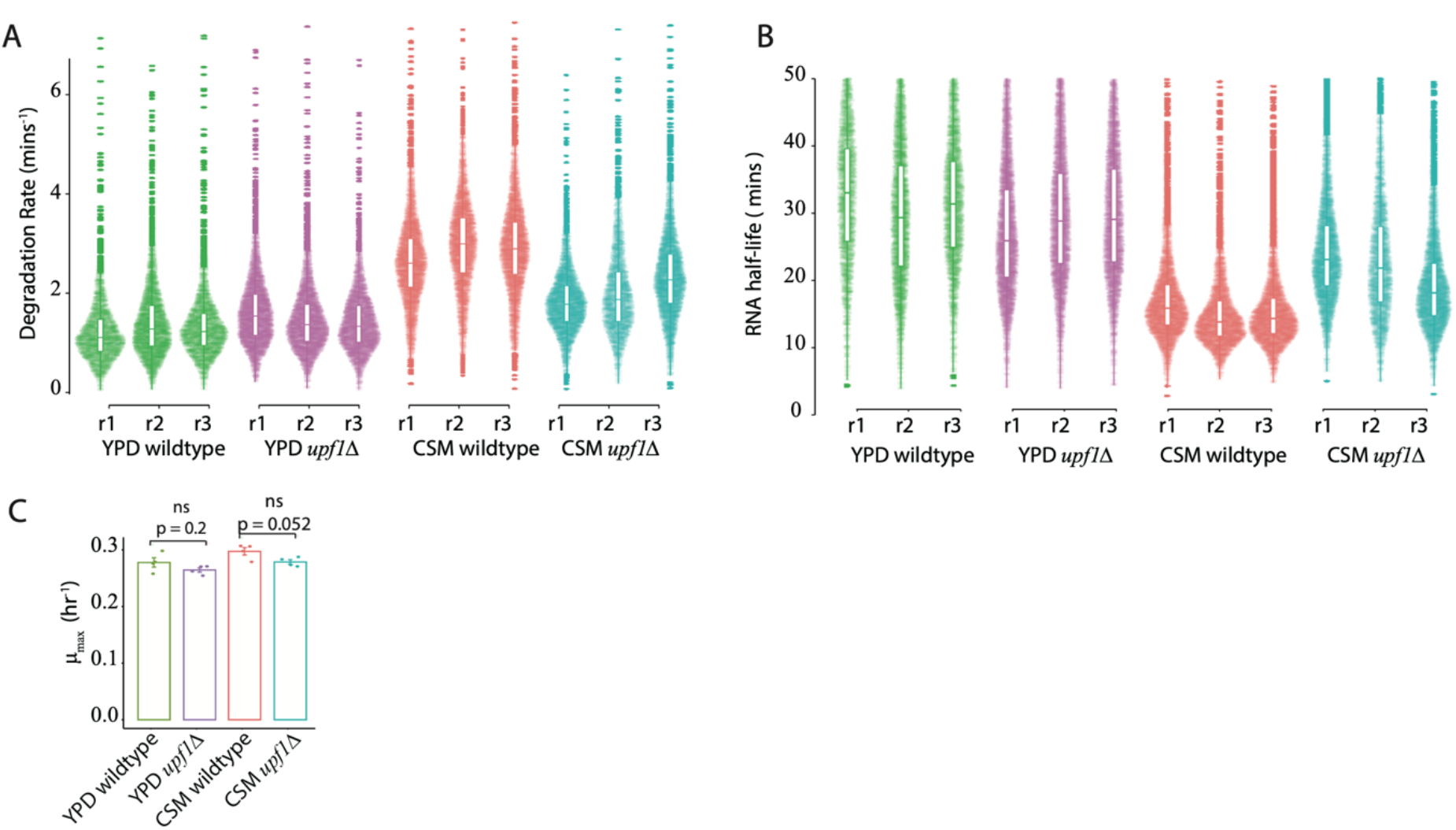
NMD and frameshifting-dependent decay in low nutrients conditions, Related to Figure 3. (A) Violin plot comparing the degradation rate (mins ^-1^) between wildtype (BY4741) and NMD mutant (*upf1Δ*) both in YPD and CSM across all three biological replicates. Only reads containing at least 1 T>C conversions were considered as labelled reads. Only coding mRNA with at least 20 total reads were considered for RNA degradation calculations. (B) as (A), but displayed as RNA half-life (in mins). (C) Bar plot comparing the growth rate (h ^-1^) between wildtype (BY4741) and NMD mutant (*upf1Δ*) both in YPD and CSM. Each condition consists of four biological replicates. Statistical analysis was performed using two-sided t tests.

**Figure S4.**
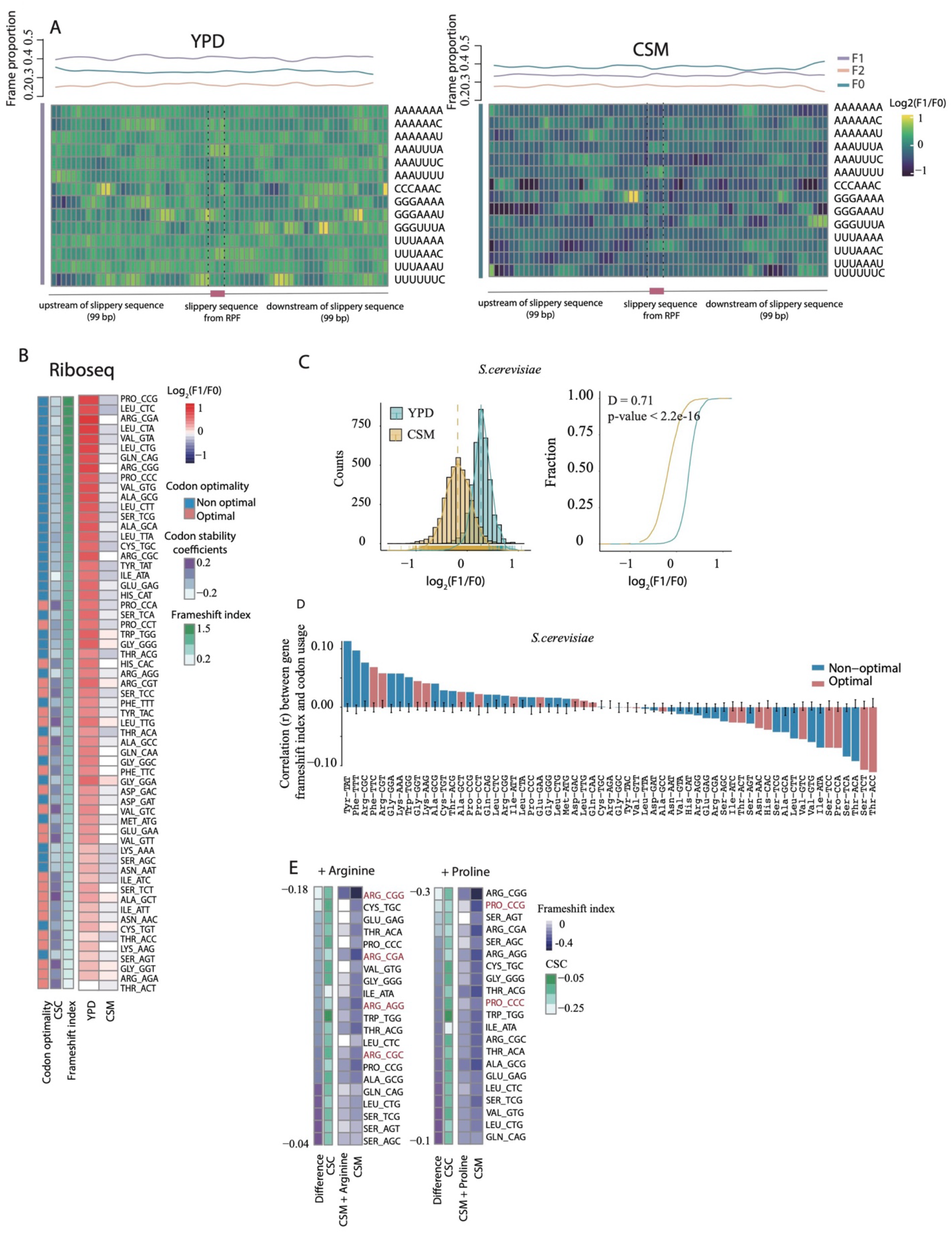
Codon optimality controls out-of-frame mRNA decay, Related to Figure 4. (A) Heatmap comparing 5PSeq coverage across three frames (F0, F1 and F2) with respect to slippery sequences in YPD (left) and CSM (right). Slippery sequence regarding to programmed ribosomal frameshift (PRF) were obtained from PRFdb^22^ (Supplementary Data 4). The average proportion of each frame are showed with sliding window of 3 nucleotides extended to 99 nt both upstream and downstream of slippery sequence. (B) Heatmap comparing ribosome profiling coverage at log_2_ ratio of -15 (corresponding to F1) and -16 (corresponding to F0) relative to the A-site of codons in YPD (Data from Santos, *et.al* ^62^) and CSM (from this study). The codons ordered by the differences of frameshift index between YPD and CSM. Codon stability coefficients (CSC) and Codon optimality were obtained from Presnyak *et al*.^10^. (C) Density plot comparing gene-specific frameshift index distributions between *S. cerevisiae* growing in YPD and in CSM using 5PSeq (left). A cumulative fraction plot with the frameshift index distribution is shown (right). Statistical analysis was performed using Kolmogorov–Smirnov test with P-value and D as effect size. (D) Correlation coefficients (Pearson) between frameshift index and codon compositions among frameshift genes (log_2_(F1/F0_) control_ > 0.2 and log_2_(F1/F0) _treatment_ < -0.2) in *S. cerevisiae*. Optimal codons and non-optimal codons are showed in red and blue, respectively. (E) Heatmaps comparing the change in codon frameshift index for the top 30% of codons, when supplemented with arginine (from 50 to 85.6 mg/L) (left) and proline (from 0 to 85.6 mg/L) (right) as compared to CSM alone. The codons are ordered by the differences of frameshift index between CSM and the respective amino acid additions.

**Figure S5.**
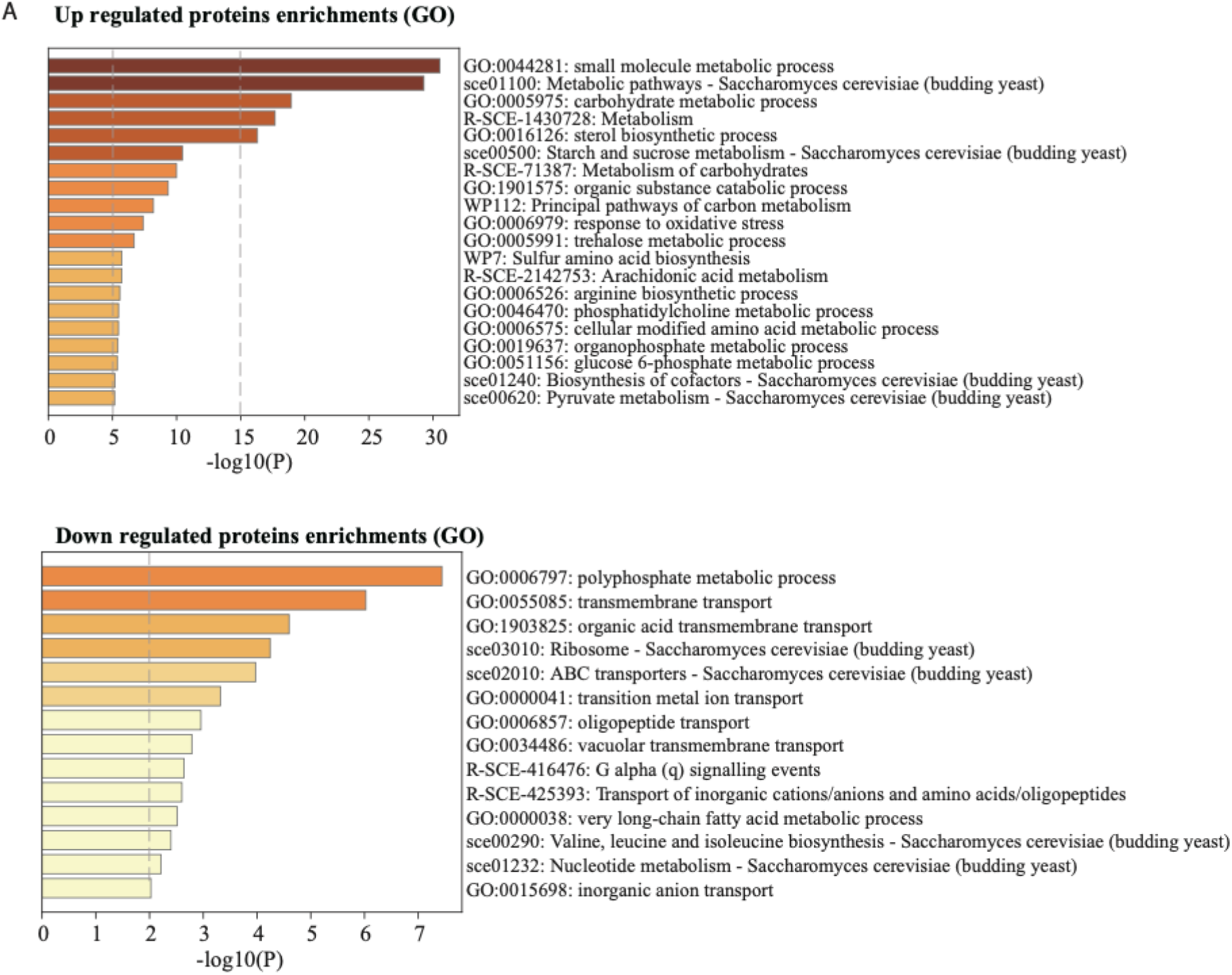
Proteomics change upon long-term amino acid limitation, Related to Figure 5. (A) Gene ontology terms for up-regulated and down-regulated protein abundance. Only statistically significant enrichments are showed (p-adj < 0.05).

**Figure S6.**
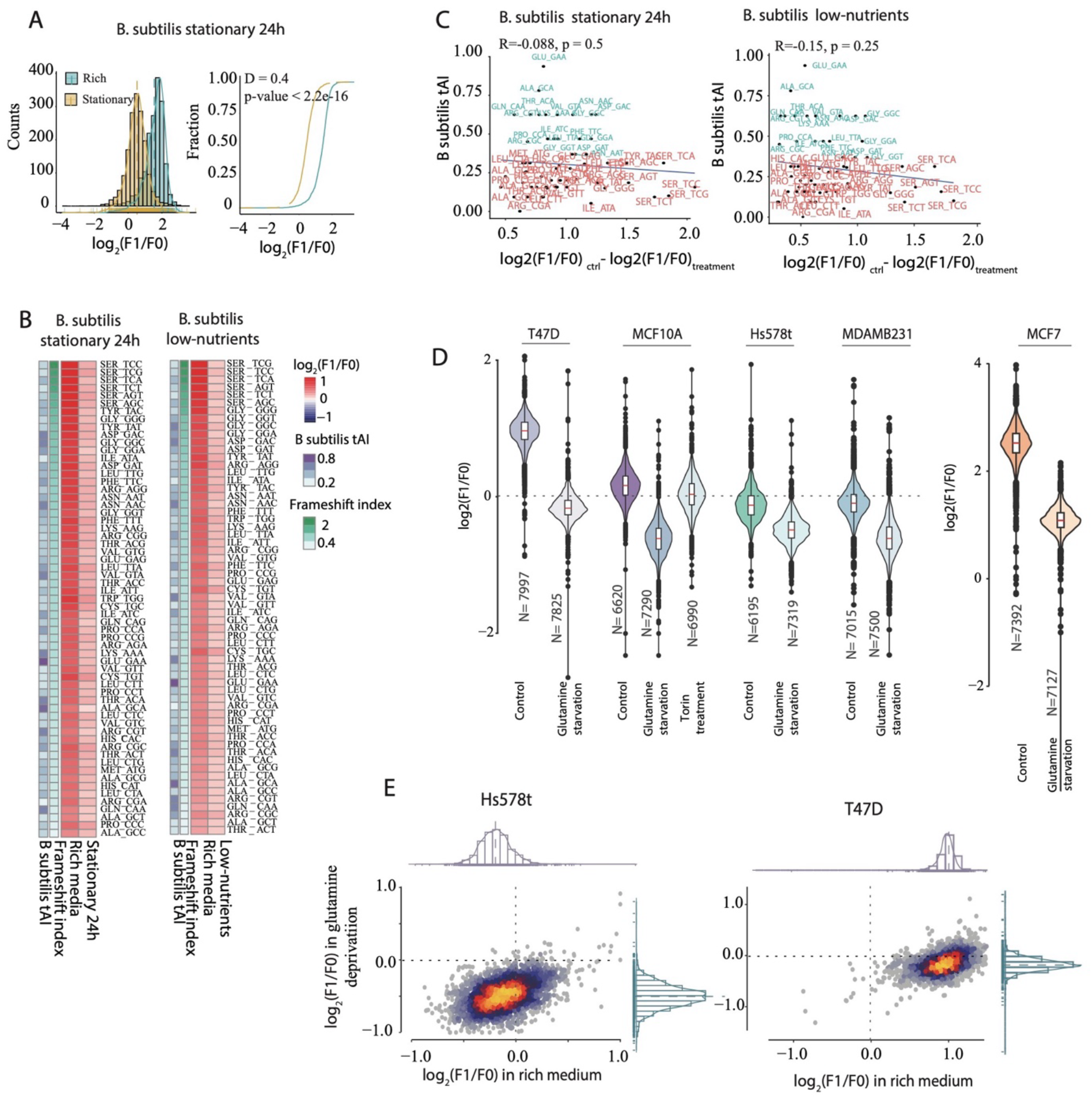
Out-of-frame co-translational decay of mRNA is evolutionary conserved, Related to Figure 6. (A) Density plot comparing frameshift index distributions between *B. subtilis* growing in stationary for 24h and in rich medium (LB). Data obtained from Huch, *et.al*.^11^ (B) Heatmap comparing 5PSeq coverage at log_2_ ratio of -14 (corresponding to F1) and -15 (corresponding to F0) relative to the A-site codons for *B. subtilis* growing in LB and in stationary phase for 24h (left). *B. subtilis* growing in poor minimal growth medium (minimal growth medium) (Details in Method section). (C) Scatterplot comparing frameshift index (x-axis) and tRNA adaptation index (y-axis) for *B. subtilis* growing in stationary for 24h (upper) and *B. subtilis* growing in poor minimal growth medium (below). The correlation between the two datasets was evaluated using Spearman correlation analysis. Optimal codons and non-optimal codons are represented in red and blue, respectively. (D) Frameshift index distributions across different breast cancer cell lines MCF10A, MCF7, T47D, MDA-MB-231, and Hs578T measured by Ribosome profiling. Data obtained from Loayza-Puch, *et.al*^36^. A generalised frameshift can be observed for all cell lies after glutamine deprivation. The red line in box plot indicates the median gene frameshift index. (E) Scatter plot comparing the frameshift index (log2(F1/F0)) of individual genes in rich DMEM medium (x-axis) and glutamine-deprived conditions (y-axis). Namely those genes with log2(F1/F0) < 0 in each condition. The upper panel in corresponds to the Hs578t cell line, while the lower panel corresponds to the T-47D cell line.

## SUPPLEMENTARY TABLES

**Table S1, Related to Figure 1**. Growth medium composition in CSM and SC.

**Table S2, Related to Figure 2**. Frameshift index in YPD, CSM and stress conditions.

**Table S3, Related to Figure 3**. RNA half-life measured by SLAM-Seq in wildtype and *upf1Δ*.

**Table S4, Related to Figure 4**. Codon frameshift index measurement in 5PSeq and Riboseq

**Table S5, Related to Figure 5**. MS measured canonical protein abundance.

**Table S6, Related to Figure 6**. Gene-specific frameshift index in bacteria and human.

